# Distributed resonant coupling enables high efficiency power transfer to mm-scale bioelectronics

**DOI:** 10.64898/2026.07.16.739014

**Authors:** Sunghoon Rho, Garrett Knuf, Atharv Naik, KunHyung Roh, Kecheng Wang, Seeva Lakshmi Cherukuri, Akash Pai, Sawson Taheri, Constantine Sideris, Siddharth R. Krishnan

**Author notes:** Corresponding author E-mail Address (Professor Siddharth Krishnan).

## Abstract

Miniaturized implantable bioelectronics offer breakthrough potential in disease treatment, biomarker monitoring, and physiological sensing. However, wireless power transfer (WPT) remains a central limitation for millimeter-scale devices, as scaling down the receiver size rapidly decreases coupling efficiency due to tissue attenuation, low quality-factors, reduced mutual inductance, and limited tolerance to spatial and angular displacement. Here, we introduce distributed resonant coupling (DRC), a 3-coil WPT paradigm which enables the receiver (Rx) to actively participate in a strongly coupled resonance, transforming the Rx from a passive energy harvester into an active participant in a strongly coupled regime. By co-designing transmitters (Tx), resonators (Rs), and mm-scale receiver coils (Rx) as fully coupled systems, DRC exhibits simulated maximum power levels of 66%, measured received power transfer efficiencies of ∼56%, and end-to-end DC power transfer efficiencies of 42%, delivering >420 mW to loads at 1 W of transmitted power while maintaining robust performance across a range of practically relevant orientations and tissue media. Systematic theoretical and experimental efforts establish core design rules for DRC systems enabling operation without specialized tuning integrated circuits or components. To illustrate the capabilities of DRC in practical applications *in vivo*, we demonstrate three technologies that capture a broad application space in bioelectronics: implant localization, rapid wireless battery charging, and ultraminiaturized drug delivery devices compatible with particulate drug formulations. Taken together, these results suggest operational capabilities across a wide range of angular tolerances (up to 60°), tissue depths and dielectric and scattering media.

## Main

Power requirements place practical limitations on the size and lifetime of most implantable medical devices (1–3). Implant technologies such as cardiac pacemakers, deep brain and nerve stimulators and cochlear implants achieve long lifetimes via large primary batteries that have limitations in packaging, safety, size and implantation site. Wireless power transfer (WPT) technologies based on near-field (4,5), mid-field (6,7) far-field (8), magnetoelectric (9,10) or ultrasound (11) energy transfer typically exhibit power transfer efficiencies (PTE) of 0.1-1% (12–15), requiring large, high-power transmitters to achieve practical levels of power beyond superficial skin depths. Embodiments of millimeter-scale receivers are based on passive harvesting paradigms (16,17), which limits WPT systems to low-power electronics applications such as sensing and communication, but well below practical thresholds required to achieve reliable rapid battery charging, deep-tissue applications or novel high-power biomedical systems such as drug delivery devices (2,3). Strongly coupled inductive power transfer regimes approach WPT efficiencies of ∼ 40-50 % over distances of ∼ 2.25 m (18) , based on transmitter and receiver coils that have diameters of >60 cm, and cannot be used in biomedical applications. These coil sizes and geometries are governed by constraints in receiver dimensions that exhibit rapidly decreasing PTE with size due to low quality factors, dielectric tissue loss, and increased sensitivity to spatial displacement and angular misalignment (19,20).

Here, we report distributed resonant coupling (DRC), a WPT architecture that can achieve >40% PTE, to mm-scale, implantable receivers. Systematic theoretical investigations reveal the active participation of 3-coil architectures in resonance that yield strong coupling with moderate Q-factors (∼68) in the loaded quality factor of the coupled resonant system. Detailed experimental investigations yield a set of design rules in establishing coupling and loss rates to maintain operation in strongly coupled regions across a wide range of practically relevant conditions, including angular tolerances up to ∼60° degrees, and through several centimeters of biological tissue. These results are achieved without any automated tuning circuits, or customized integrated circuit design. We demonstrate 3 new biomedical technologies enabled by strongly coupled mm-scale systems. We first demonstrate rapid charging of implantable batteries, with currents of up to 78 mA, allowing for complete battery charge cycles in <1 hour, a capability which can shift implantable device powering paradigms from primary to secondary cells. We then illustrate a novel, complementary capability in return-loss based localization through thick tissue layers, as a mechanism to spatially locate implants without any visual cues. Finally, we demonstrate an ultraminiaturized battery-free implantable drug delivery that is compatible with indefinitely stable, solid formulation drugs, with operational modes that rely on thermally actuating shape-memory alloy (SMA) metals *in vivo*. This collection of results spans new physical insights in mm-scale 3-coil resonance, technological advancements in wireless power transfer, and enabling demonstrations that motivate new medical technologies that can exploit high-efficiency power transfer for a range of clinical needs.

### Distributed resonant coupling for millimeter-scale wireless power transfer

DRC comprises a 3-coil topology in which the transmitter (Tx), an intermediate resonator (Rs), and a millimeter-scale receiver (Rx) are co-designed to form a single strongly coupled resonant system across three antennas (**Fig. 1a**, Extended Data Fig. 1a). Tx and Rs are planar coils, with diameters of 3.6 and 4.6 cm, respectively while Rx, the implantable component is ∼3.5 mm x 4.5 mm, and has a total mass of 0.12 g, comparable to a grain of rice (**Fig. 1a** top right). The return loss S_11_ spectrum of the lumped Tx-Rs-Rx DRC system reveals shared resonant modes two characteristic peaks (21,22) at the eigenfrequencies of the combined system (**Fig. 1b**). In contrast, the spectrum for Tx alone exhibits a single shallow minimum (return loss >−4 dB) and no peak splitting. These measured spectra yield several key system parameters, including the loaded quality factor of the coupled resonant system (Q ∼ 68), the coupling rate (K ∼ 1.21 × 10⁶ s⁻¹), and the loss rate (Γ ∼ 6.22 × 10⁵ s⁻¹), averaged across four different receiver antennas. Critically, the resulting average coupling-to-loss ratio, K/Γ, is 1.95 at the system resonance frequency of 13.56 MHz (**Fig. 1b** and Extended Data Fig. 1c), well above the strong-coupling threshold of K/Γ = 1 (21,22) . Numerical simulations (**Fig. 1c**) reveal the presence of strong coupling only when all 3 coils actively participate in resonant energy transfer, a property of this system that distinguishes it from traditional strong-coupling regimes (18) . As such, the magnetic flux stored as circulating current in the intermediate Rs is coupled into the Rx, thereby enabling several important emergent properties such as measured PTE of up to 56% in optimized load conditions, approaching simulated maximums of 66% (**Fig. 1d**). WPT systems typically exhibit monotonic dependences between Rx size and PTE, a tradeoff that has limited scaling to implantable, mm-scale regimes (5,6,9,13,15,23–32). Active participation across 3 coils based on DRC modes addresses this tradeoff, as shown in **Fig. 1e** and Supplementary Fig. 1. where a simple, computed ratio between PTE and Rx cross sectional area as (19) serves as one figure of merit and total received power serves as a second. These scaling advances suggest a broad set of applications and capabilities in implantable medical technologies that are currently limited by power budgets (**Fig. 1f**).

**Fig 1.**
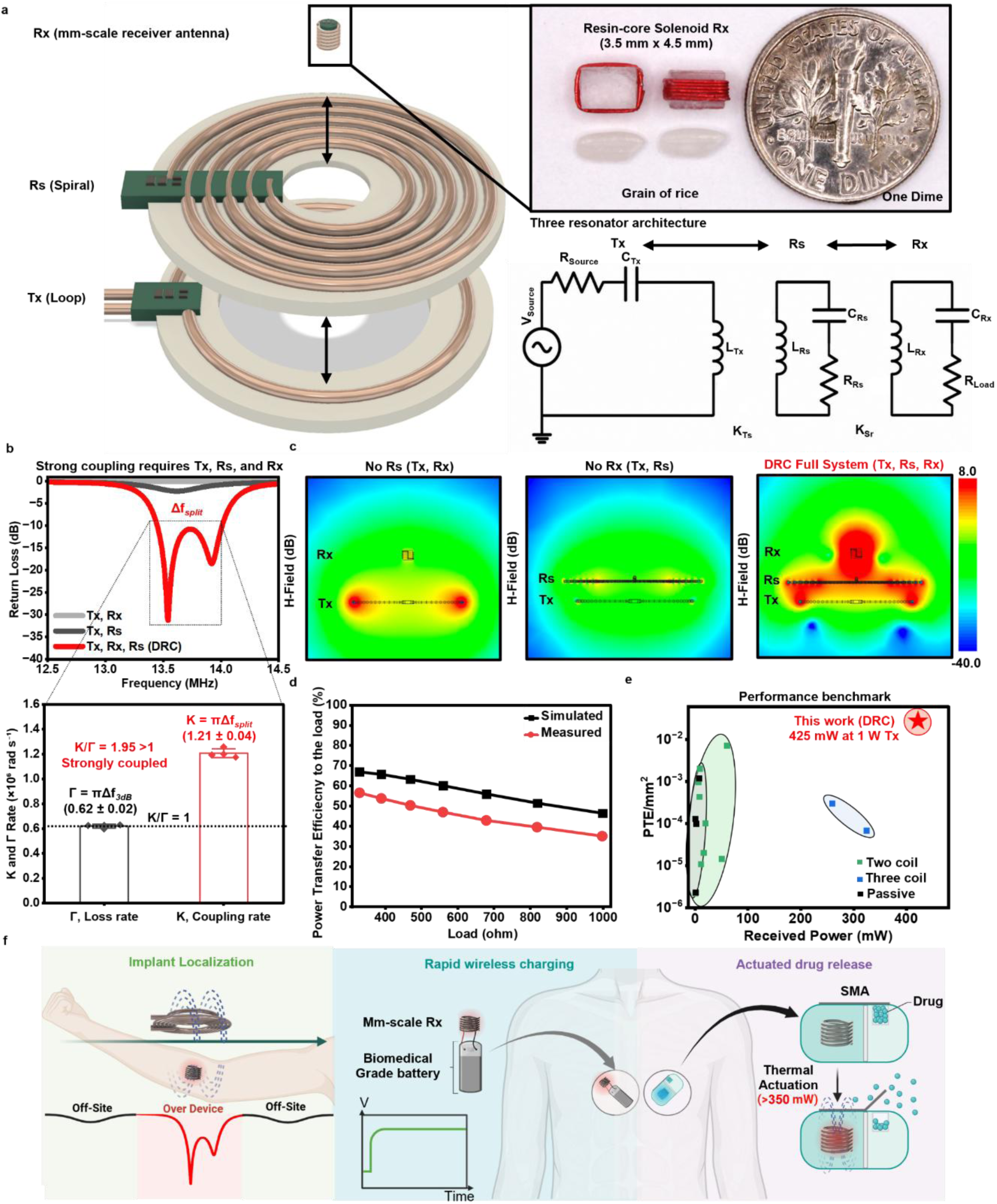
Overview of distributed resonant coupling (DRC) for high-power wireless bioelectronics. **a**, Three-dimensional illustration of the DRC architecture consisting of a transmitter loop (Tx), an intermediate spiral resonator (Rs), and a millimeter-scale receiver (Rx). Photographs of the fabricated receiver alongside a grain of rice and a US dime highlight its miniature dimensions. The equivalent circuit representation of the three-resonator architecture is shown**. b,** Measured return loss (S_11_) of the complete DRC system and calculated K/Γ for 4 different receiver antennas (Bars represent mean ± SD, n = 4 measurements). Simulated magnetic-field (H-field) distributions for the full architecture and control configurations without Rs and Rx in **c**, Strong coupling emerges only when the Tx, Rs, and millimeter-scale Rx simultaneously participate in the resonant mode, resulting in mode splitting and efficient wireless power transfer. **d,** Comparison of measured and simulated results across different load conditions. **e,** Benchmark comparison of wireless power transfer systems showing delivered power as a function of receiver size, highlighting the high received power enabled by DRC. **f,** Representative high-power bioelectronic functions enabled by DRC, including implant localization, wireless charging of miniaturized electronics, and wireless actuation for on-demand therapeutic delivery.

### Strongly coupled resonant modes enable high-power transfer to mm-scale receivers

Systematic experimental studies highlight key requirements and operational parameters of integrated DRC architectures (**Fig. 2a**) across a range of settings relevant to practical biomedical implant design. The S_11_ peak splitting, a characteristic signature of strongly coupled systems is consistently observed across laterally displaced regions (left, center, right of the inner loop of Rs) within the resonator aperture (**Fig. 2b**), suggesting that the DRC conditions are not artifacts arising from diode nonlinearity or parasitic circuit components. Implant technologies often demand regulated DC loads and comparative studies between DRC circuits with and without rectification and load components (diodes, capacitors, resistors) exhibit equivalent peak-splitting behavior (Extended Data Fig. 1a,b) suggesting that the strongly coupled mode is an intrinsic property of DRC and independent of parasitic behaviors of these components, with a simple theoretical framework for these behaviors appearing in Supplemental Note S1. Experimental evaluations establish end-to-end wireless power transfer from the transmitter to the load, as a measure of usable DC output power (**Fig. 2c**). The Tx–Rs distance was set to 5 mm based on the optimization results shown in Supplementary Fig. 2. At an input power of 1 W, compatible with many batteries, the DRC system delivers 424 [±40] mW of usable DC power to the load at a distance of 1 cm, corresponding to an end-to-end PTE of 42.4 [±4]%. This value was measured across the fully integrated receiver, including the rectifier, storage capacitor, and load, thereby evaluating the overall end-to-end efficiency of the DRC system. These power measurements are substantially higher than those of 2-coil configurations such as Tx-Rx (∼1.2 mW) and Rs-Rx (∼56.3 mW) under the same input power conditions (**Fig. 2c**). Notably, the system presented more than 100 mW of delivered power at transmission distances approaching 2.5 cm, corresponding to depths relevant for many subcutaneous and intramuscular implants (33). Additionally, increasing Tx power is a simple pathway to higher total received DC power; for example, increasing input power from 1 W to 2 W results in received DC power of ∼665 mW, suggesting possibilities in scaling DRC architecture to higher power levels (**Fig. 2d**). Safety considerations in specific absorption rate (SAR) represent an important, practical requirement of our system. Numerical simulations predict a peak 1g averaged SAR of 1.176 W/kg under the maximum power-transfer condition, with the receiver embedded in tissue which is below the FCC SAR limit 1.6 W/kg (SAR1g) (Extended Fig. 1h and Supplementary Fig. 3) (34). The antenna temperature increased in air because of the low thermal conductivity of air, encapsulation with biomedical clear resin effectively controlled this heating, limiting the temperature increase to less than 5 °C over (up to 60 seconds) (Extended Data Fig. 2j and Supplementary Fig. 4).

**Fig 2.**
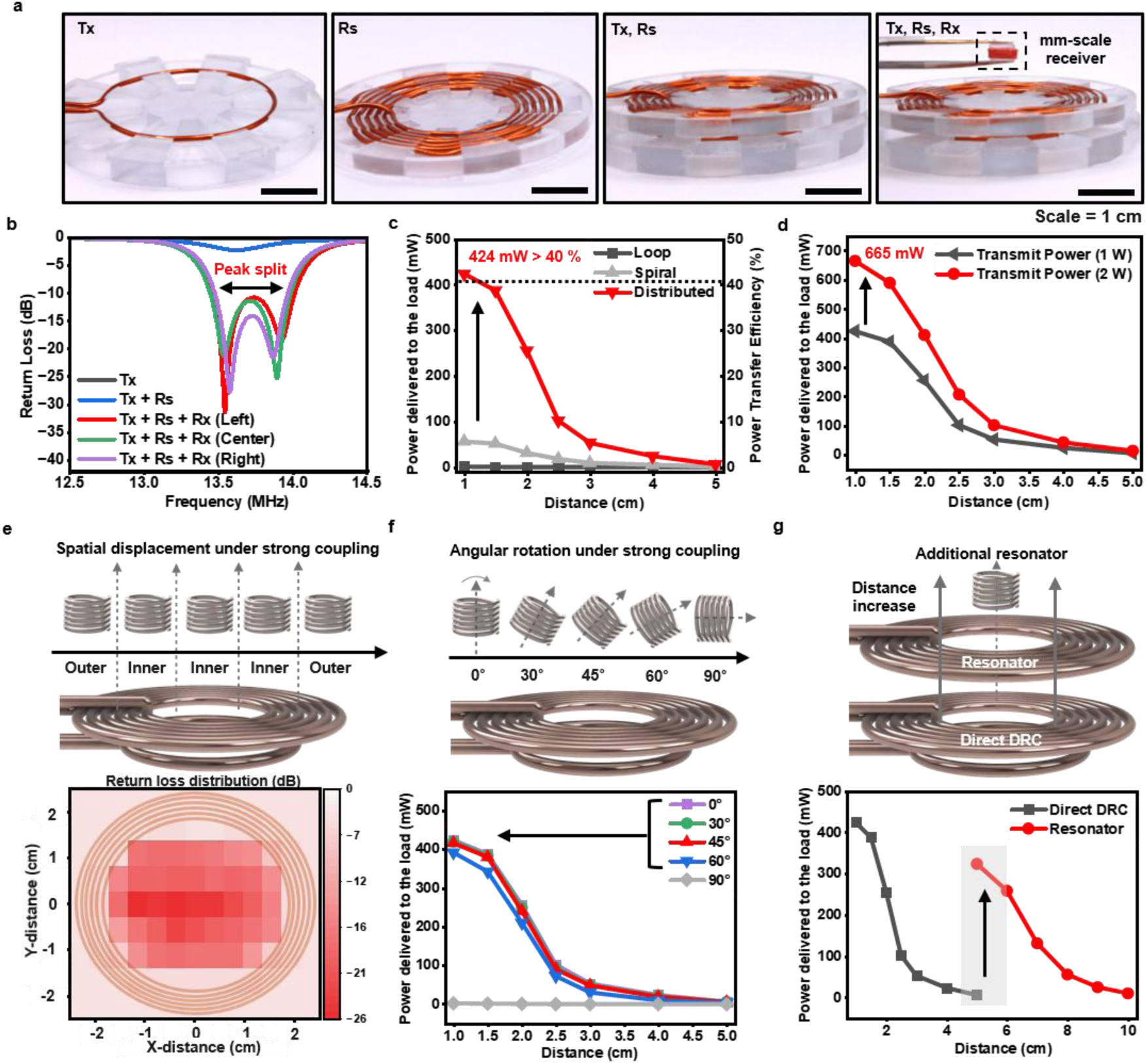
Physics of distributed resonant coupling for high-power millimeter-scale bioelectronics. **a**, Distributed resonant coupling (DRC) architecture comprising a parallel-resonant transmitter (Tx), an intermediate spiral resonator (Rs), and a millimetric receiver (Rx). The co-designed resonant elements collectively form a strongly coupled wireless power transfer system (scale bar = 10 mm). **b,** Measured return loss (S_11_) of the Tx antenna positioned at the left, center, and right sides of the inner loop of the spiral resonator (Rs). The frequency-split peaks at a coupling distance of 1 cm presents the strongly coupled regime. **c,** Received power delivered to an optimized 1,000 Ω load at a coupling distance of 1 cm for different transmitter. Only the DRC enables high-power transfer to millimeter-scale receivers, achieving 424 mW of received power. **d,** Power scaling characteristics of the DRC system. Received power increases linearly with transmitter power, reaching 665 mW as the transmitted power is increased from 1 W to 2 W. **e, ,** Spatial displacement tolerance of the DRC system. High power delivery is observed across substantial receiver displacements within the transmitter loop area, demonstrating spatial robustness. **f,** Angular robustness of the DRC architecture under strong coupling. Efficient power transfer is maintained across a wide range of receiver orientations and coupling distances upto 60**°**. **g,** Scalability of the DRC architecture through the addition of a secondary resonator. An additional resonator at a distance of 5 cm extends the strongly coupled regime and provides an additional 300 mW of received power, highlighting the potential of DRC for long-range wireless power delivery.

Power transfer tolerances to spatial and angular misalignments represent an important parameter for many bioelectronic devices, particularly in contexts where implants cannot be visually assessed. Strongly coupled power transfer regimes offer high levels of uniformity in power transfer across resonator apertures, as shown by measurements of return loss across the resonator (**Fig. 2e**). Additionally, tolerance to angular misalignment represents a key feature of these systems: a simple analysis that accounts for the cosine distribution of coupled magnetic flux suggests that strong coupling (K/Γ>1) can be maintained up to an angle of cos^-1^ (1/2.0) ∼ 60° (**Fig. 2f** and Supplementary video 1). Experiments and HFSS simulations validate these concepts, suggesting power transfer efficiencies that are conserved up to ∼ 60° in functional DRC systems. Full-wave simulations reproduced the measured self-impedances of the three antennas, and the simulated angle- and distance-dependent power transfer closely agreed with the experimental measurements (Extended Fig. 1f-g). Additionally, compared with planar coils of a similar receiver areas (∼200 mm²) (12,15), the proposed DRC architecture delivered approximately fourfold higher received power under both angular misalignment and increased transmission distance (Supplementary Fig. 5). Finally, we note strategies in 4-coil based design to enhance the range of DRC-based power transfer to up to 10 cm, corresponding to deep-brain and other deep tissue regions (35), based on the addition of a passive resonator (**Fig. 2g** and Supplementary video 2 and 3). This resonator enhances the received power by approximately 300 mW. These antennas represent simple, hand-wound coils without specialized tuning or matching circuitry. In this context, we evaluated the reproducibility of DRC architectures via fixed transmitter–resonator (Tx–Rs) pairs with five independently fabricated receivers 399.7 ± 10.5 mW (mean ± standard error of the mean, N = 5). The measured power across these pairs exhibited a standard deviation of less than 5%. Additional experiments deployed fixed receivers with 12 independently fabricated Tx–Rs pairs, all of which produced consistent wireless power-transfer performance 417.4 ± 5.2 mW (mean ± standard error of the mean, N = 12) (Extended Data Fig. 1e, and Supplementary video 4). In addition, the DRC architecture exhibits a threshold behavior that enables the millimeter-scale receiver to actively participate in the strongly coupled resonant mode (Extended Data Fig. 2a–g). Owing to the relatively weak load dependence of these systems, the measured end-to-end PTE remained above 40% for load resistances up to 1 kΩ before decreasing at 2 kΩ (Extended Data Fig. 2h). Additional experiments characterize the effects of the Rx design, including wire dimensions, the number of turns, and receiver load impedance under high-resistance loading conditions (Extended Data Fig. 2i). Furthermore, the presence of a large iron object adjacent to or directly above the Rx reduced the received power by only ∼15%, demonstrating the robustness of the proposed system to nearby metallic interference (Supplementary Fig. 6 and Supplementary video 5). Implantation of the receiver in the subcutaneous and intraperitoneal spaces of a mice cadaver resulted in only ∼10% attenuation of the received power (Supplementary Fig. 7).

### Localization and charging in freely moving mice

*In vivo* transcutaneous power transfer represents an important set of capabilities with utility across a broad range of implants (3,36). A key challenge in this context is poor angular or spatial alignment in the absence of visual cues (37). Strong coupling is evidenced by peak splitting in the S_11_ spectrum, where both split resonances exhibit return-loss values of approximately −10 dB. This insight suggests capabilities in localization of implantable receivers through several millimeters of opaque tissue and other optically dense media based on simple, readily available hardware such as handheld vector network analyzers (VNAs) (**Fig. 3a**). Measurements over tissue locations adjacent to the implanted device with the fully integrated battery, PMIC, and antenna, in living mice show shallow return-loss spectra (S_11_ ∼ −3 dB) and no peak splitting, indicative of weak coupling (**Fig. 3b** and Supplementary video 6). In contrast, measurements directly over the implanted device exhibit deep return loss (S_11_ ∼ −22 dB) with characteristic peak splitting, whereas measurements acquired near the device show a single deep return-loss peak (S_11_ ∼ −10 dB), enabling spatial localization of the implant (**Fig. 3b** and Supplementary video 6). As such, this approach enables localization of implanted receivers with integrated antenna circuits through simple tissue scanning and return-loss heatmap reconstruction in cadavers (**Fig. 3c** and Supplementary video 7). Experiments with intraperitoneal implants in mouse cadavers demonstrate these concepts, with spatially resolved return-loss heat maps identifying implant location, at depths of ∼ 10 mm without any visual cues. Benchtop experiments using tissue phantoms, including dense chicken breast tissue, suggest that these concepts can be extended up to ∼10 mm of tissue beneath the implant and 10 mm above it, corresponding to implantation depths of approximately 1-3 cm (Supplementary Fig. 8-9). Localization represents a complementary capability to battery charging.

**Fig 3.**
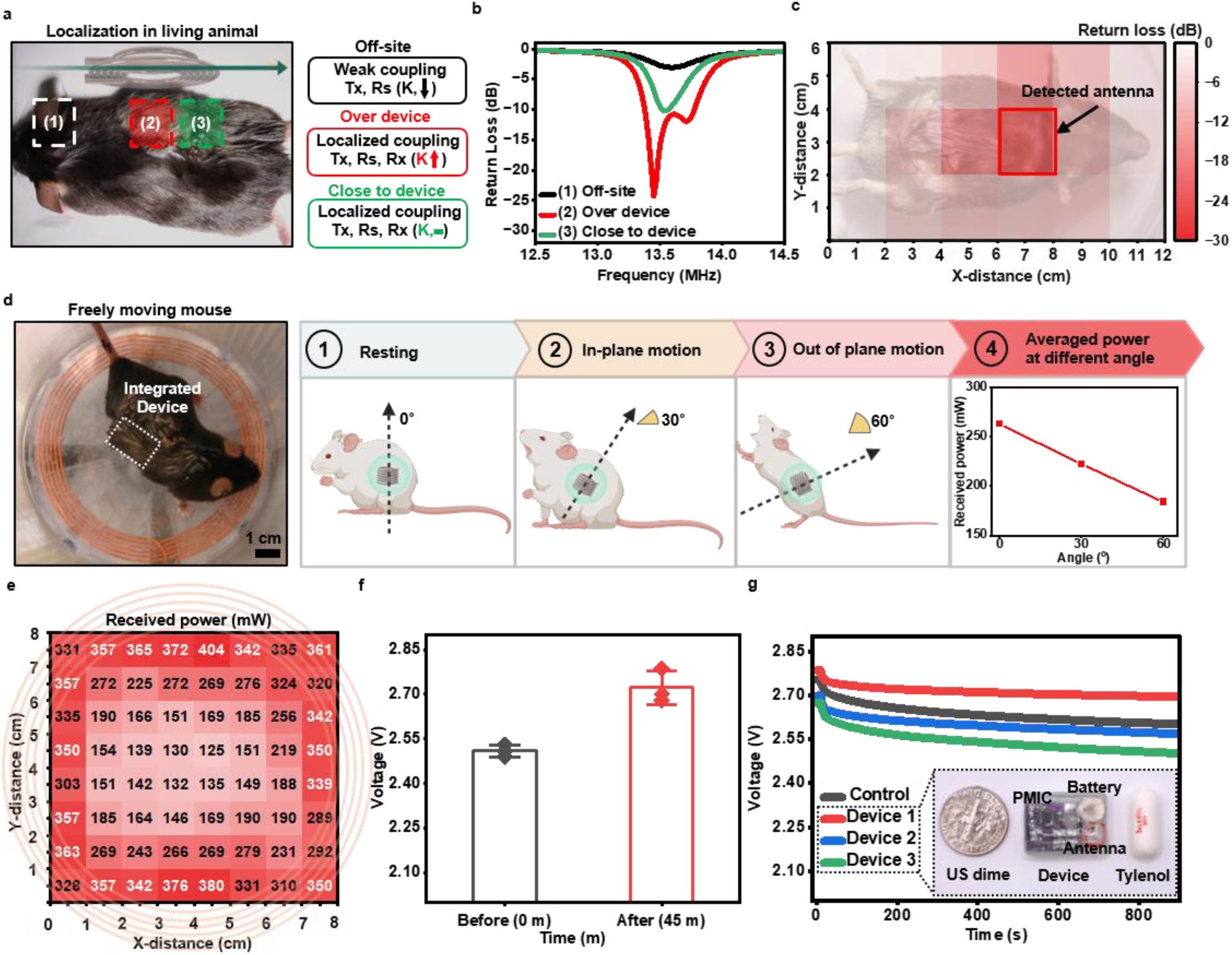
Localization and wireless power transfer for freely moving implanted bioelectronics. **a**, Schematic illustration and photograph of implant localization using distributed resonant coupling (DRC) in a live mouse. The implanted receiver actively participates in the strongly coupled resonant mode, producing characteristic frequency-split peaks in the transmitter return-loss (S_11_) spectrum when measured directly over the implant. In the absence of the implanted device, only a shallow resonance is observed. **b,** Localization of an implanted device in a live mouse using the fully integrated implant, including the PMIC, antenna, and battery. Representative S_11_ spectra are shown for measurements performed off the implant, directly over the implant, and near the implant. **c,** Measured return-loss maps demonstrate spatial localization of an implanted receiver without the PMIC circuit in a cadaver model, with the deepest return loss observed directly above the implanted antenna. **d,** Wireless power transfer to a freely moving mouse with an integrated implantable device. A photograph of a freely moving mouse positioned above the transmitting antenna is shown, with the implanted device indicated by the white box. Representative schematics of device orientations during freely moving operation, including resting, in-plane, and out-of-plane orientations, together with the corresponding average received power. **e,** Received power distribution across the transmitting area with the device in the resting (0°) orientation. **f,** Measured battery voltage before and after 45 min of wireless charging. The biomedical-grade rechargeable 3 V, 5.8 mAh lithium coin-cell batteries increased from 2.51 ± 0.02 V to 2.72 ± 0.06 V (mean ± SD, n = 3). **g,** Battery discharge curves of the three wirelessly charged devices. Devices with similar initial voltages but without wireless charging were used as a control. All three wirelessly charged devices exhibited comparable discharge properties, indicating successful battery charging. A photograph of the fully integrated device is also shown alongside a U.S. dime and a Tylenol tablet for size comparison.

DRC platform further enabled rapid wireless charging using a power-management integrated circuit (PMIC) and a 3.7 V, 40 mAh Li-ion battery with the handheld Tx-Rs. Charging currents of up to 78 mA were achieved at a transmitter power of 2 W and approximately 60 mA at 1 W (Supplementary Fig. 10). More than 10 mA of charging current was maintained at a transmission distance of 4 cm. The charged batteries were subsequently discharged using a potentiostat, confirming successful wireless energy storage and repeatable charging performance. For charging in awake, freely moving mice, we designed a large (∼10 cm diameter) transmitter incorporated into a 3D-printed housing. Studies in murine cadavers in these systems (Supplementary Fig. 11) demonstrated rapid charging of 40 mAh lithium polymer batteries in <1 hour. These custom housing systems were constructed with Rx-Rs distances of 1 cm and enabled continuous charging across a practical range of positional misalignments of 30° (received power 222 mW) to 60°(184 mW) (Supplementary Fig. 12). A complete distribution of received power transfer profiles across the Tx aperture appears in **Fig. 3e**. *In vivo* charging in freely moving animals involved the use of a miniaturized, biomedical grade, 3V 5.8 mAh battery (Supplementary video 8). Following an initial complete discharge *ex vivo* and 45 minutes of charging, we observed an increase in measured battery voltage and standard discharge curves consistent with those of unused batteries (**Fig. 3g**). The measured temperatures of freely moving mice at 0, 20, and 40 min were 30.53 ± 0.51 °C, 30.70 ± 0.75 °C, and 31.30 ± 0.36 °C, respectively (mean ± SD, n = 3; Supplementary Fig. 13).

### Wireless therapeutic actuation enabled by distributed resonant coupling

Drug delivery platforms for complex biologic drugs represent an emergent class of bioelectronic technologies (38–40). Traditional drug delivery systems based on passive-release biomaterials, or mechanically actuated pumps have limitations in control and drug stability, respectively (41–43). These latter challenges in stability frustrate the development of active delivery systems for many peptide and protein based biologic drugs that exhibit short shelf-lives in aqueous forms, frustrating long-term storage (12,44). Particulate versions of these drugs such as lyophilized formulations offer indefinite stability but are challenging to deliver. Recent technologies have demonstrated novel actuation mechanisms based on the ablation of thin-gold films (41) or the mechanical actuation of shape-memory alloy (SMA) actuators (12,42,45–48). Both of these technologies require large actuation currents (0.1-1A) that are supported by large batteries. Here, we demonstrate the first entirely battery-free drug active drug delivery device for particulate drug formulations, enabled by DRC. These systems suggest pathways to combine miniaturization and shelf-stability, without the need for battery charging and based on validated concepts (3,49) in long shelf-life particulate forms of drugs.

Our design (**Fig. 4a**) comprises a DRC receiver antenna, printed circuit board (PCB) supporting resistive joule heater and tuning and rectification components, an SMA actuator, a drug reservoir, and encapsulation integrated into a millimeter-scale package with a size comparable to that of oral tablets. Briefly, the DRC Rx drives a current, *I*, through the heating strip comprised of surface-mounted resistors in series with a total resistance *R* that are mechanically coupled to the SMA actuator. Local joule heating, given by *I^2^R* driven entirely by wirelessly harvested power then provides a stimulus for local SMA shape transformation based on phase change, that can open a reservoir and release a particulate drug (**Fig. 4b**). Previous studies have shown that SMA actuation requires a minimum power of >350 mW (12), and the integrated board facilitates heat dissipation from the resistor (**Fig. 4c and Supplementary Fig. 14**) to achieve this. The fully integrated device was first evaluated in water, representing a highly lossy dielectric environment, with a high thermal conductivity (kwater ∼ 0.6 W/m-K) relative to that of dry air (kair ∼ 0.01 W/m-K). Upon wireless power delivery, the SMA actuator successfully opened the drug reservoir in water, resulting in the time-dependent diffusion of a model dye-based test drug system, Rhodamine B, into the surrounding solution (**Fig. 4d** and Supplementary video 9). To evaluate *in vivo* performance, the device was loaded with lyophilized fluorescent model drugs based on Cy7 and implanted into subcutaneous regions in live mice. Following 30s of wireless power transfer, the harvested energy triggered SMA actuation, resulting in successful opening of the implanted device (**Fig. 4e**). *In vivo* opening was observed via *in vivo* imaging systems (IVIS) based on the appearance and continuous expansion of a fluorescent diffusion front for up to 20 min following wireless activation. This diffusion was not observed in the control group, in which the devices were implanted but not actuated at the same transmitter power of 2 W (**Fig. 4f**). A total of six mice were used (three control and three actuated). All three actuated devices successfully opened their SMAs following wireless activation. The IVIS dye diffusion image shown is representative of one of the three actuated mice. Together, these results demonstrate that DRC delivers sufficient wireless power to drive battery-free mechanical actuation and controlled particulate molecule release *in vivo*. By enabling therapeutically relevant power delivery to millimeter-scale implants, DRC establishes a practical foundation for next-generation implantable bioelectronics requiring large power budgets.

**Fig 4.**
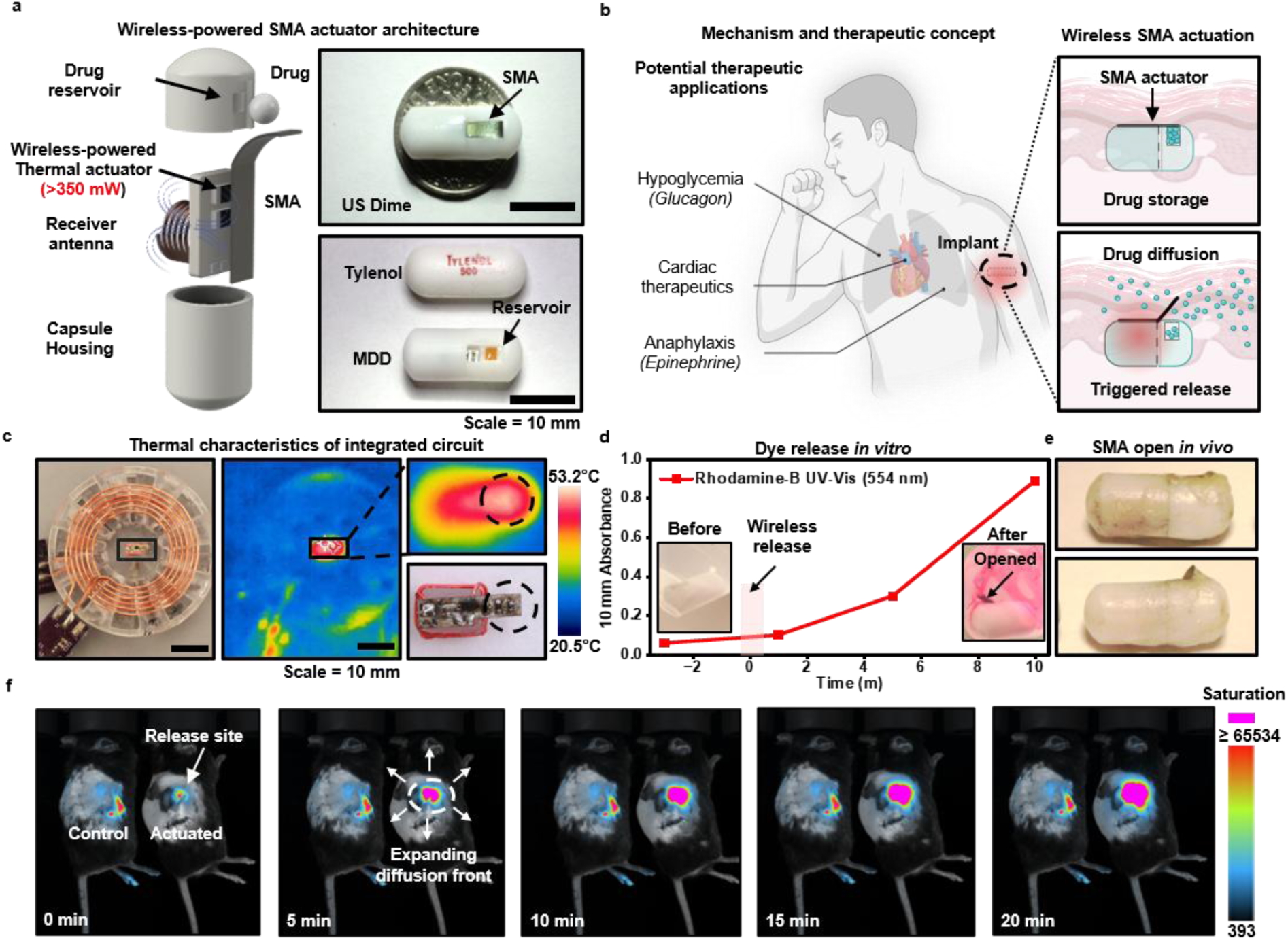
Miniaturized drug delivery (MDD) platform powered by distributed resonant coupling for wireless therapeutic actuation. **a**, Exploded-view illustration of the miniaturized drug delivery (MDD) device, showing the wireless SMA actuator, drug reservoir, and triggered-release mechanism. Photographs of the MDD device are shown alongside a US dime and a Tylenol tablet for size comparison. b, Schematic illustration of potential therapeutic applications enabled by DRC-powered implantable devices, including treatments for hypoglycemia, cardiac disorders, and anaphylaxis. c, Wirelessly powered device for thermal dissipation. Representative near-infrared thermal images are shown in the middle. A magnified view of the resistor region is shown at the top right, highlighting the temperature distribution at the device resistor. d, Wireless release of Rhodamine B from the implanted drug delivery device in water and the corresponding fluorescence intensity as a function of time. e, Representative explanted devices following wireless actuation in mice, confirming successful device opening and triggered drug release *in vivo*. f, Time-lapse IVIS images showing wireless drug release from the implanted MDD over time.

## Discussion

Miniaturizing high-power strongly coupled wireless power transfer systems has been limited by the large size mismatch between the transmitter and millimeter-scale receivers, where the achievable efficiency of power transfer is limited by low coupling coefficients and quality factors (18,22). Implanted biomedical antennas typically operate in regimes where receivers are function as a passive harvesters, with an associated set of drawbacks in low efficiency and alignment tolerance (50). In this work, we show that these apparent scaling limits can be overcome via the creation of shared resonant modes through co-design of three-resonator distributed resonant coupling systems. These systems allow the stored magnetic flux in passive intermediate resonator coils to be channeled into mm-scale receivers to support high-efficiency energy transfer with large end-to-end usable power budgets. DRC enables high-power delivery to mm-scale implants, supports localization measurements tissue, rapid recharging of implantable batteries *in vivo*, and battery-free drug delivery systems compatible with particulate drugs. This functionality is supported by robust performance against dielectric tissue loss and misalignment in position and orientation. We note an important limitation of this work, in that it has been validated in short, acute *in vivo* demonstrations. Additional work will evaluate long-term performance in small animal and large animal preclinical studies, with co-designed wearable or handheld transmitters for energy transfer.

These results could motivate future efforts in developing advanced reconstruction algorithms for implant localization that account for the heterogeneous anatomy of living tissue, combining amplitude, phase, frequency shift measurements. This technology can be further generalized to complex, power-intensive bioelectronics applications that currently require battery power. This is supported by the observation that a second resonator placed after the transmitter delivered an additional ∼300 mW to the receiver, suggesting that the spatial extent of strong near-field coupling can be substantially expanded across larger anatomical volumes, with potential applications in closed-loop deep-brain stimulation, cardiac pacing, and ingestible electronics (3,35,51,52).

## Materials and Methods

### Fabrication of a transmitting antenna

The transmitter loop geometry was adapted from a previously reported wireless power transfer antenna design with minor modifications to match the DRC architecture (21,22). A custom mold was fabricated using a 3D printer to guide the winding of the spiral resonator using copper wire (AWG 18, Remington, IL, USA). After fabrication, the spiral resonator was connected to a 50 Ω SMA connector, and its impedance was measured using a vector network analyzer (VNA). At 13.56 MHz, the measured impedance was 1.6 + j158.265 Ω. Because the resistive component was negligible compared with the reactive component, only the reactance was considered for resonance tuning. A parallel combination of 68 pF and 8 pF capacitors (total capacitance of 76 pF) was selected to cancel the measured inductive reactance and tune the spiral resonator to 13.56 MHz. Due to variations in the manufactured capacitor values and parasitic effects introduced by the PCB, a tuning process was performed over a capacitance range of 75 - 78 pF using combinations of a 68 pF capacitor with either an 8 pF or a 10 pF capacitor. After fabrication of the spiral resonator, the transmitter loop was characterized in the same manner. The measured loop impedance was 0.7 + j7.2 Ω at 13.56 MHz. A series tuning capacitor was selected to cancel the measured reactance and maximize the loop current at the operating frequency. For this, a 1.5 nF capacitor is connected in series to cancel the reactance. The separation between the transmitter loop and the spiral resonator was fixed at 5 mm based on the optimization results shown (Supplementary Fig. 2). CAD files for the antenna holder are provided in Supplementary Fig. 16. The transmitter has a turn-to-turn spacing of 1 mm and consists of 6 turns of 18 AWG wire (1.02 mm thickness), with a starting radius of 2.5 cm.

### Fabrication of a receiver antenna

Copper wire (AWG 26, 28, 30, 32, 34, and 36; Remington Industries, IL, USA) was used to fabricate the receiver antennas. Different wire gauges were evaluated to investigate the effect of wire thickness on coupling efficiency with various numbers of turns (2–10). A custom mold (3 mm × 4 mm × 3 mm) was fabricated using a Form 4 3D printer (Formlabs, MA, USA) to support the receiver coil during winding. After the copper wire was wound around the 3D-printed mold, the coil was secured using cyanoacrylate adhesive. Because the copper wire was enamel-insulated, the insulation was carefully removed from the wire ends using a blade before electrical connection.

### Characterization of the received power of DRC

The received power was evaluated using a fully integrated receiver circuit consisting of five components: the fabricated receiver antenna, a resonant capacitor, a rectifier, a buffer capacitor, and a load resistor. The resonant capacitor was selected by measuring the receiver impedance with a vector network analyzer (VNA) and compensating for the inductive reactance at 13.56 MHz. For example, a six-turn receiver fabricated with AWG 30 wire exhibited an impedance of 0.9 + j17 Ω, and a tuning capacitor was selected to cancel the reactive component and achieve resonance at the operating frequency. A 1 μF buffer capacitor was connected after the rectifier to reduce the output voltage ripple. Load resistances ranging from 200 Ω to 2 kΩ were used to evaluate load-dependent power transfer performance. The printed circuit boards (PCBs) were fabricated by OSH Park, Inc.. The rectified voltage across the load resistor was measured using a digital multimeter. The received power was then calculated as P=V^2^/R, where V is the measured load voltage and R is the load resistance.

### Characterization of the DRC without diode and load

To characterize the DRC system without active electronic components, a simplified receiver board was fabricated without the rectifier, buffer capacitor, or load resistor. The receiver consisted only of the antenna and the resonant capacitor, as shown in Extended Data Fig. 1a. The return loss (S_11_) of the transmitter was measured using a vector network analyzer (VNA). To capture the resonance near the operating frequency of 13.56 MHz, the frequency was swept from 12.5 to 15 MHz using 1,001 measurement points with five-point averaging. In addition, the transmission coefficient (S21) was measured by connecting the receiver board to Port 2 of the VNA through a 50 Ω SMA connector while varying the separation distance between the top of the Rs and the center of the Rx from 1 to 5 cm. The transmission coefficient was measured to evaluate the S-parameters of the unmatched circuit. The measured S-parameters were subsequently used to optimize and validate the electromagnetic model in HFSS.

### Wireless powering system

Wireless power was delivered using a commercially available modular wireless power transfer system designed for rodent experiments, consisting of a 13.56 MHz signal generator, power amplifier, and graphical user interface (Neurolux Inc., Skokie, IL, USA). The system was operated at 13.56 MHz with the continuous waveform. For DRC experiments, the commercial transmitter antenna was replaced with the custom-fabricated Tx and Rs in this work. The transmitted power was set to 1 W or 2 W, with a maximum output power of 2 W. In other words, the Neurolux system was used only as the signal generation and power amplification module, whereas wireless power delivery was performed using the custom-fabricated DRC Tx and Rs.

### Localization evaluation of the distributed resonant coupling

A NanoVNA was used to evaluate the localization capability of the DRC system in both in vitro and *ex vivo* experiments. The transmitter (Tx) and spiral resonator (Rs) were mechanically assembled into a single module. The implanted receiver board was fully integrated and encapsulated with UV-curable resin to protect the electrical circuit from moisture and liquid exposure. For the *ex vivo* localization experiments (chicken breast and cadaver), the Tx–Rs module was scanned laterally over the tissue to identify the implanted receiver. For localization in chicken breast tissue, the Tx–Rs module was scanned in both the horizontal and vertical directions to map the receiver position. To evaluate the localization resolution, the scanning step size was set to match the receiver dimensions (3.5 mm × 4.5 mm), and the measurement area was divided into grids of the same size. At each grid position, the return loss (S_11_) was measured using the NanoVNA, and the resulting values were visualized as a two-dimensional heat map.

### Battery charging of the DRC

A power management integrated circuit (PMIC, BQ25185 1-Cell) was integrated with the DRC receiver board to evaluate the wireless charging performance of the system. Because the receiver was used for battery charging, no external load resistor was connected. The receiver circuit consisted of the receiver antenna, resonant capacitor, rectifier, buffer capacitor, and PMIC. A 10 μF buffer capacitor was used to reduce the rectified voltage ripple and to provide a stable input to the PMIC during charging. The PMIC was configured according to the manufacturer’s datasheet. A 43 kΩ resistor connected to the ILIM/VSET pin was used to program the input current limit, while a 3 kΩ resistor connected to the ISET pin was used to set the battery charging current. The TS/MR pin was connected through a 10 kΩ resistor according to the manufacturer’s recommended configuration. For the handheld version (high-power efficiency), the same Tx–Rs was used to charge the integrated board with the antenna and PMIC circuit shown in Supplementary Fig. 10. For the freely moving experiment, a larger Rs was used to provide sufficient space for the mice to move. For this, the same Tx dimensions were used, but two turns were applied to increase the mutual coupling between the relatively small Tx and the larger Rs. We used a 3V, 5 mAh rechargeable biomedical grade secondary battery, chosen for its size compatibility with freely moving mice (ML621-TZ1, FDK America, Inc.)

### Miniaturization of the drug delivery device

The Tylenol-sized, single-dose device was fabricated from three main components; Housing, SMA, and integrated board with antenna. SMA sheets, machined by electrical discharge machining to dimensions of 2.3 mm × 15 mm × 0.1 mm and programmed with a transition temperature of 43 °C (Kellogg’s Research Laboratories), were used as described previously (12). Sulfo-Cyanine7 carboxylic acid was dissolved in PBS at a concentration of 6 µg µl⁻¹. To prepare a near-indefinite dry formulation, the dye solution was loaded into a 10 mm³ drug reservoir and lyophilized. The antenna was soldered onto the printed circuit board (PCB) together with 0201-size capacitors (710 pF, comprising 560 pF and 150 pF). A buffer capacitor, rectifier diode, and a 260 Ω resistor (implemented using two 130 Ω resistors in series) were integrated into the circuit. To improve heat dissipation, thermal paste was applied to the top surface of the resistor, followed by stacking of the shape-memory alloy (SMA) actuator. After device integration, heat dissipation was characterized (Supplementary Fig. 14). Device localization in 1% agarose gel is shown in Supplementary Fig. 15. The CAD files for the encapsulation are provided in Supplementary Fig. 16.

### Device injection in mice and IVIS imaging for SMA actuation

All procedures were supervised by the Stanford Institutional Animal Care and Use Committee under protocol number 34823. C57BL/6J mice (approximately 20 weeks old, ∼35 g) were used for device implantation. All animals were anesthetized with 3% isoflurane in O₂, and the dorsal surgical area was shaved and sterilized with povidone-iodine (Betadine) followed by 70% ethanol. A 2-cm incision was made in the upper right dorsal skin, and a subcutaneous pocket was created using blunt forceps. The device was inserted into the pocket, and the incision was closed with 5-0 nylon sutures. All procedures were performed in accordance with institutional guidelines for perioperative care, anesthesia, and pain management. One day after implantation, *in vivo* fluorescence imaging was performed using an IVIS imaging system (Scintica Newton FT-900). A cyanine dye (Cy7) was used to visualize wireless dye release *in vivo*. Images were acquired using the Cy7 emission filter with a fixed exposure time of 40 ms. The excitation intensity and aperture settings were adjusted according to the fluorescence intensity of the implanted device in the mice real-time (Supplementary Fig. 17). Before wireless actuation, 100 μL of sterile saline was injected into the implantation site to facilitate dye diffusion. Immediately after the saline injection, wireless power was delivered through the skin for 30 s at a transmitting power of 2 W.

### Electromagnetic Modelling and Simulation in HFSS

Full-wave electromagnetic simulations were conducted in Ansys HFSS to further validate the empirical performance of the distributed resonant coupling. The simulated Tx, Rs, and Rx geometries were precisely modelled to mirror the physical dimensions of the fabricated AWG copper coils, including wire gauge, conductor spacing, number of turns, and loop diameter. To match the loading environment at the receiver, a dielectric resin core (εr = 3.0) was placed at the center of the receiver coil with the dimension of 3.5 mm x 4.5 mm. Each coil structure was excited via differential lumped ports isolated by a Perfect Electric Conductor (PEC) bridge. The multi-coil assembly was centered within a 60cm cubic air box (10x the largest coil dimension) which was terminated with a Finite Element-Boundary Integral (FE-BI) boundary condition. The simulated self-impedances for the Tx, Rs, and Rx were measured to be 0.027 + 9.22j, 0.193 + 140.3j, and 0.211 + 14.31j at 13.56 MHz, respectively, which verifies close matching to the physical Tx, Rs, and Rx. Since the spacing between coils heavily affects DRC behavior, the following geometries were parameterized: Tx-Resonator distance (2 mm to 14 mm), Resonator-Rx distance (0.5 cm to 5 cm), Rx tilt angle (0° to 90° around its center of mass). For each geometric variation, adaptive spatial meshing was iteratively performed at the 13.56 MHz solution frequency until the maximum delta S-parameter magnitude between successive passes converged below a 2% threshold. To characterize the spectral response and verify the presence of multi-coil frequency splitting modes, a frequency sweep was performed from 12 MHz to 15 MHz using a discrete frequency sweep configuration.

The multi-port S-parameter network generated by HFSS was exported into an Ansys Circuit Design co-simulation environment. The circuit schematic is shown in Extended Fig. 1a., and the corresponding capacitor values selected were optimized to resonate each coil at 13.56 MHz. A 1W, 50 Ω sinusoidal power source drives the series capacitor connected to the Tx coil to excite the network. At the receiving end, a port connects in parallel with the LC tank acting as the load. The load nominally assumes a value of 1 kΩ, but can be adjusted to observe the load dependence on the system. To extract the system S-parameters, a linear network analysis was conducted, sweeping the frequency of the sinusoidal input power source from 12 MHz to 15 MHz. The power transfer efficiency (PTE) was defined as the ratio of active power delivered to the terminating load resistor relative to the maximum available power from the power source, calculated as PTE (%) = |S21|^2^ x 100. This formulation penalizes the efficiency calculation for source-side reflection losses (S_11_ mismatches), which more realistically captures the realizable power transfer efficiency in miniaturized systems.

To complement the network analysis, full-wave magnetic field distributions (H-field) were plotted in the 3D domain. The field plots were generated by mapping the steady-state amplitude and phase data derived from the circuit co-simulation directly back onto the 3D solver ports and finite element mesh. This approach ensures that the visualized magnetic field contours accurately represent the true, physically loaded operating state of the multi-coil system. At last, the SAR was calculated using a tissue density of 1g cm⁻³, following the standard FCC evaluation procedure. SAR was evaluated using a finite element method (FEM) solver in accordance with IEEE 62704-4.

## Contribution

S.R. and S.R.K. conceptualized the physics and device architecture and designed the overall study. S.R., K.H.R., and A.N. designed, optimized, and assembled the devices. S.R. and S.L.C. developed the CAD design and device architecture. G.K, C.S, A.P., K.H.R. performed electromagnetic simulations and modeling. K.W., and S.R. conducted the *in vivo* experiments. S.L.C. assisted with drug-device formulation and preparation. S.R. and S.R.K. analyzed and organized the data. S.R. and S.R.K. wrote the manuscript with input from all authors. S.R.K. supervised the study. All authors read and commented on the manuscript.

## Acknowledgements

S.R.K gratefully acknowledges funding from the Leona and Harry M. Helmsley Charitable Trust and the National Institutes of Bioengineering and Bioimaging (NIH-NIBIB). A.N. gratefully acknowledges funding from The Lewis M. and Barbara C. Terman Graduate Fellowship. We also thank the Tianqiao and Chrissy Chen Ideation and Prototyping Laboratory (Chen IPL) at Stanford University for providing equipment and for their collaboration.

---

**Extended Data Fig 1.**
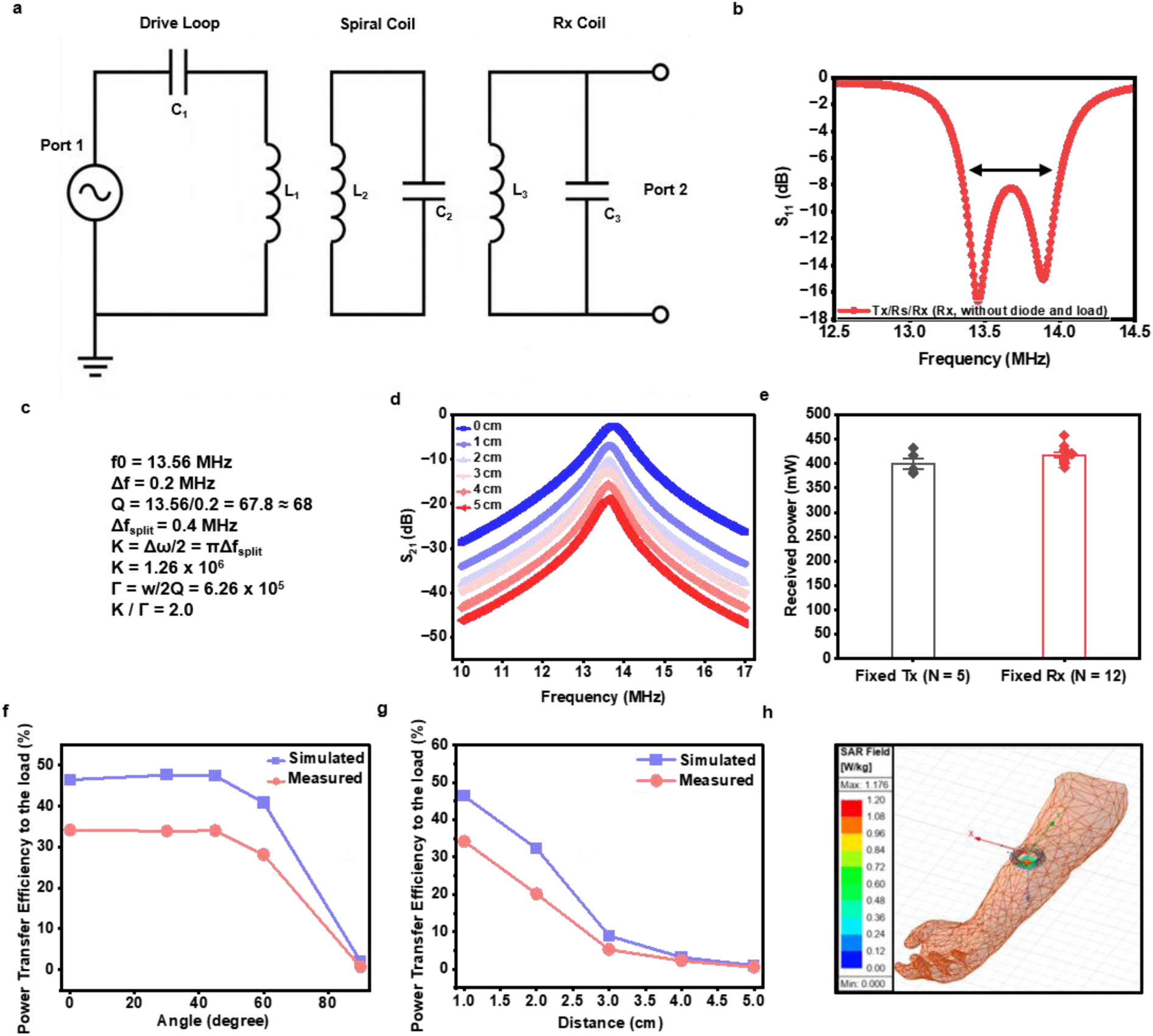
Characterization of distributed resonant coupling for millimeter-scale wireless power transfer. **a,** Equivalent circuit model of the distributed resonant coupling (DRC) system consisting of the transmitter (Tx), spiral resonator (Rs), and receiver (Rx). For S-parameter characterization, the receiver includes only the resonant capacitor, without the diode or load. **b,** Measured return loss (S_11_) of the Tx, Rs, and Rx.**c,** Calculated K/Γ based on the measured return-loss spectra. **d,** Measured transmission coefficient (S_21_) as a function of transfer distance, with the receiver consisting only of the resonant capacitor (without the diode or load). **e,** Reproducibility evaluation using the same Tx and Rs with five different Rx devices, and the same Rx with twelve different Tx–Rs pairs. **f,** Simulated and measured power transfer efficiency (PTE) of the DRC system as a function of angular misalignment. **g,** Simulated and measured PTE as a function of transfer distance. h, Specific absorption rate (SAR) simulations performed in HFSS using the same coil geometry as the fabricated devices.

**Extended Data Fig 2.**
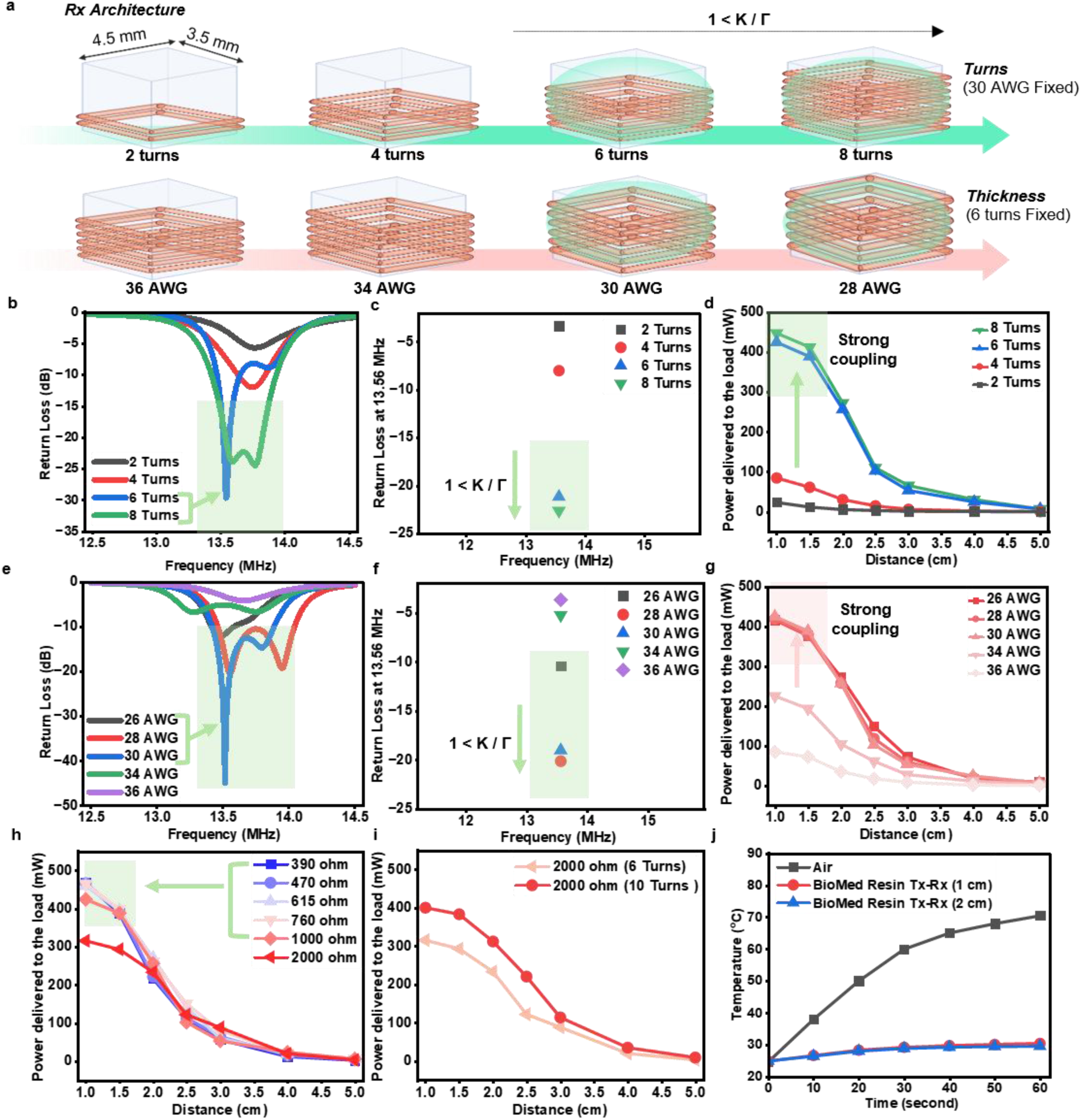
Optimization of the distributed resonant receiver for millimeter-scale wireless power transfer. **a.** Geometric illustration of the millimeter-scale receiver antenna designed to achieve strong coupling. **b.** Analysis of return loss (S_11_) for varying coil turns (2, 4, 6, and 8), with **c**, highlighting the return loss specifically at 13.56 MHz. **d.** Power delivered to the load (mW) as a function of the number of turns. **e.** Coil thickness on return loss for a 6-turn configuration, with **f** showing the return loss at 13.56 MHz. **g.** Received power as a function of coil thickness. **h.** Power delivery evaluation across various load resistances, ranging from 390 ohm to 2000 ohm. **i.** Enhancement of delivered power levels using increased turns with a 2000 ohm load. **j.** Temperature changes over time for the device in air and coated with biomedical resin at distances of 1 cm and 2 cm.

