## Supplementary Figures 1-17 for "Distributed resonant coupling enables high efficiency power transfer to mm-scale bioelectronics"

### Supplementary Information

| Order | Papers | Receiver Area ($mm^2$) | Tx Power (W) | Rx Power | PTE (efficiency) | PTE/mm2 |
|---|---|---|---|---|---|---|
| 1 | Kiani et al | 1256.00 | ~0.7 W | 260 mW | 37.00% | 0.0295% |
| 2 | Ye et al | 380.00 | | 9.2 mW (max output) | 75.40% | 0.1984% |
| 3 | Li et al | 70.90 | 0.12 W | 60 mW (102 mW max) | 50.00% | 0.7052% |
| 4 | Krishnan et al | 200.00 | 12 W | 20 mW | 2.00% | 0.0100% |
| 5 | Rho et al | 3.14 | 6 W | 8 mW | 0.13% | 0.0424% |
| 6 | Rho et al | 3.14 | 2 W | 6 mW | 0.30% | 0.0955% |
| 7 | Ouyang et al | 256.00 | 4 W | 11 mW | 0.28% | 0.0011% |
| 8 | Yokoi et al | 400.00 | 2 W | 16 mW | 0.80% | 0.0020% |
| 9 | Krino et al | 77.00 | 3 W | 0.5 mW | 0.02% | 0.0002% |
| 10 | Cai et al | 875.00 | 4 W | 50 mW | 1.25% | 0.0014% |
| 11 | Burton et al | 800.00 | 6 W | 325 mW | 5.42% | 0.0068% |
| 12 | Bansal et al | 3.00 | 2 W | 7 mW | 0.35% | 0.1167% |
| 13 | Chen et al | 44.40 | 6 W | 1.17 mW | 0.01% | 0.0002% |
| 14 | Woods et al | 22.50 | 10 W | 2.2 mW | 0.22% | 0.0098% |
| 15 | John S Ho | 3.14 | 0.5 W | 0.2 mW | 0.04% | 0.0127% |
| 16 | Ours | 15.75 | 1 W | 424 mW | 42.40% | 2.6921% |

**Supplementary Fig 1.** Detailed data underlying the Pareto plot in Fig. 1e. The referenced studies are compared in terms of receiver (Rx) coil area, received power, and power transfer efficiency (PTE)

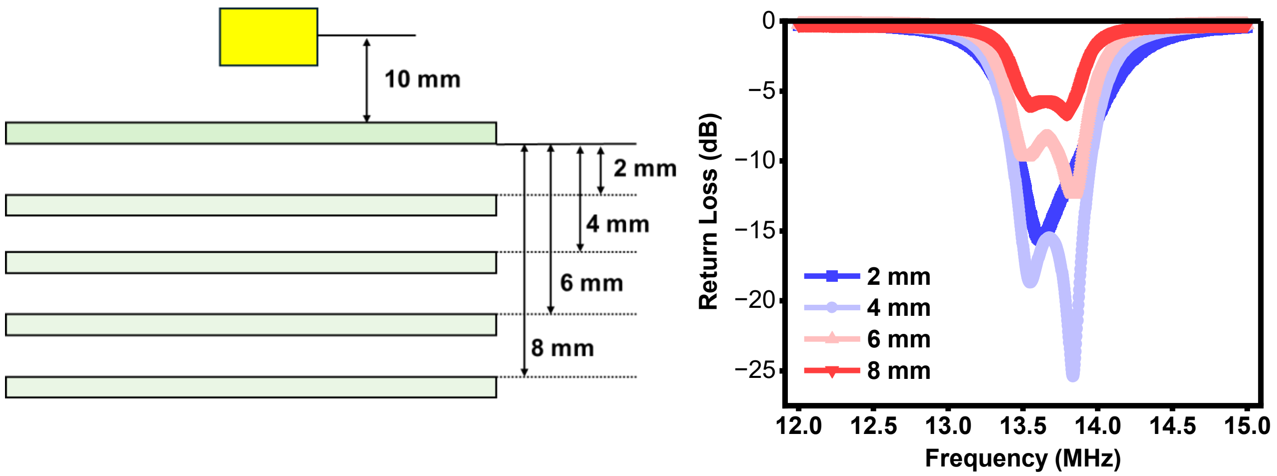


**Supplementary Fig 2.** $S_{11}$ sweeps were measured between the spiral resonator and loop at separation distances ranging from 2 to 8 mm, while keeping the distance between the top of the spiral resonator and the receiver fixed. With the distance from the top of the spiral resonator to the center of the receiver fixed at 10 mm, $S_{11}$ remained below −10 dB for spiral - loop separations of 4 - 6 mm.

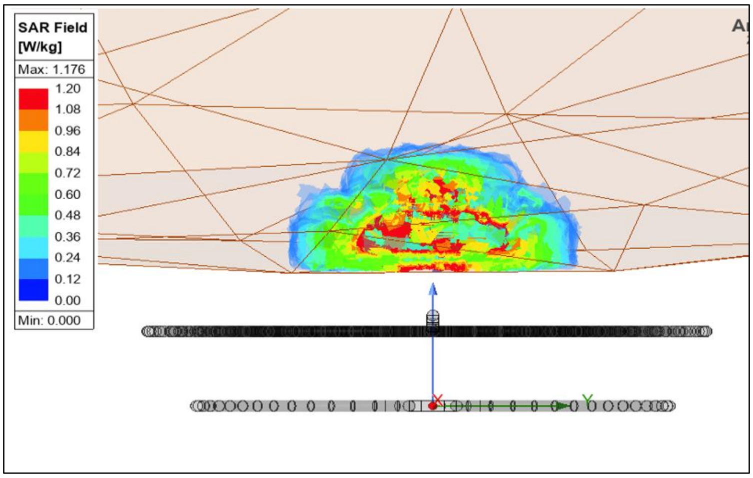


**Supplementary Fig 3.** Zoomed-in simulated specific absorption rate (SAR) distribution at a separation distance of 0.5 cm from the spiral coil in the arm tissue. The simulation was performed using a tissue density of 1 g/cm³ with 1 g spatial averaging, consistent with the standard SAR evaluation under federal communications commission (FCC).

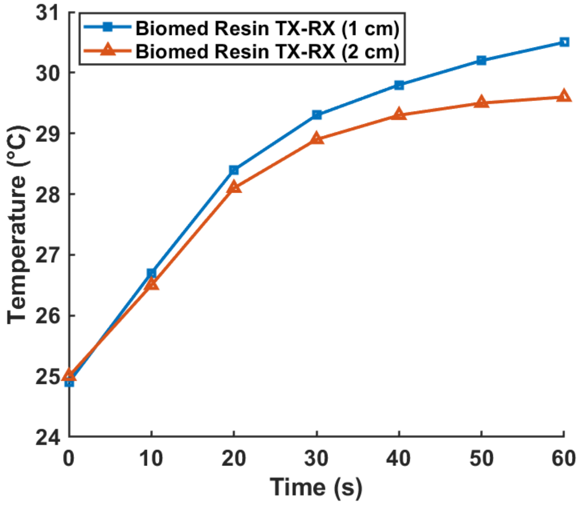


| Type | Thermal Conductivity |
| --- | --- |
| Air | 0.024 W/(m.k) |
| Biocompatible Resin | 0.25 ~ 0.35 W/(m.k) |

**Supplementary Fig 4. Temperature characterization of the Biomed Resin-coated antenna.** Zoomed-in temperature profiles of the Biomed Resin-coated antenna are shown for transmitter (Tx)–receiver (Rx) distances ranging from 1 to 2 cm. A table summarizing the thermal conductivities of air and Biomed Resin is also provided.

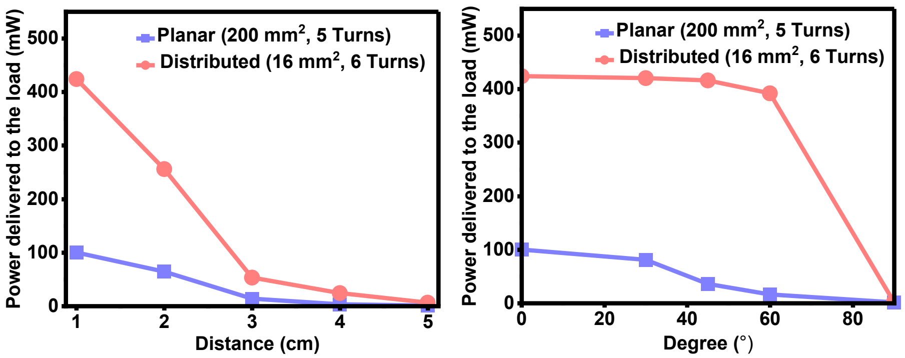


**Supplementary Fig 5. Comparison of the received power as a function of distance and angle between conventional planar resonant coupling and distributed resonant coupling (DRC).** Using the same transmitter (Tx) and spiral resonator (Rs) with different receiver (Rx) configurations, DRC achieves approximately fourfold higher received power than the conventional planar receiver.

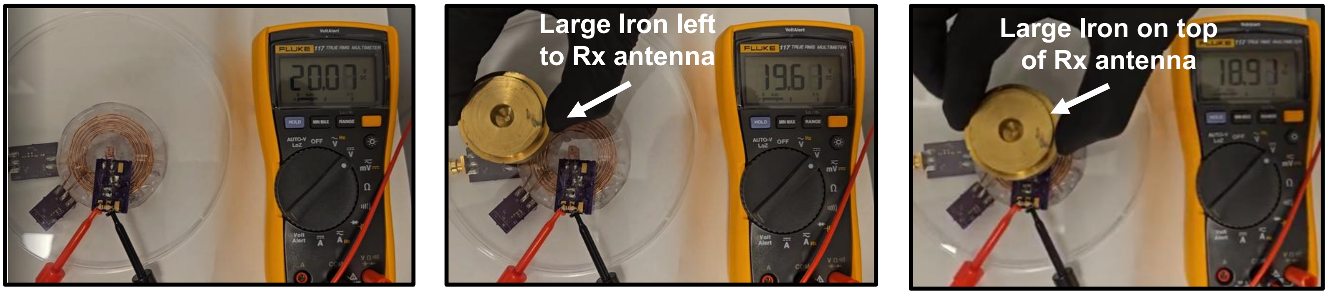


**Supplementary Fig 6. Effect of an iron object placed near the DRC system.** Placement of a large iron object near and above the receiver (Rx) reduced the received power; however, the measured output power remained above 300 mW, with measured voltages of 19.61 V and 18.91 V across a 1 kΩ load. The voltage across the 1 kΩ load on the receiver board was measured using a multimeter. This end-to-end measurement directly accounts for parasitic losses, printed circuit board (PCB) resistance, matching network efficiency, and antenna losses.

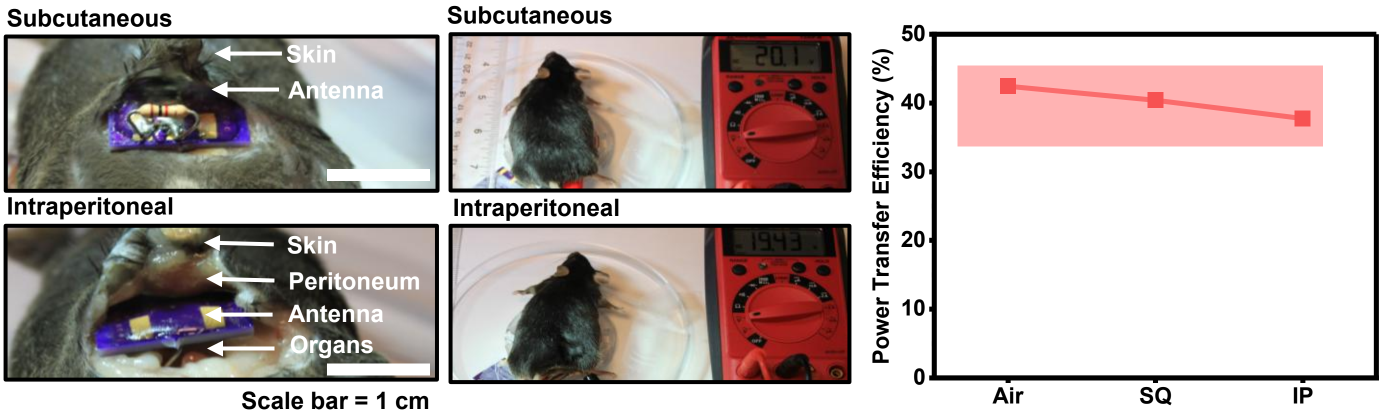


**Supplementary Fig 7. Received power in the subcutaneous and intraperitoneal spaces of cadaver mice.** Received power was measured using a millimeter-scale receiver implanted in either the subcutaneous or intraperitoneal space under the same experimental conditions. A comparison of the received power measured in air, the subcutaneous (SQ) space, and the intraperitoneal (IP) space is presented.

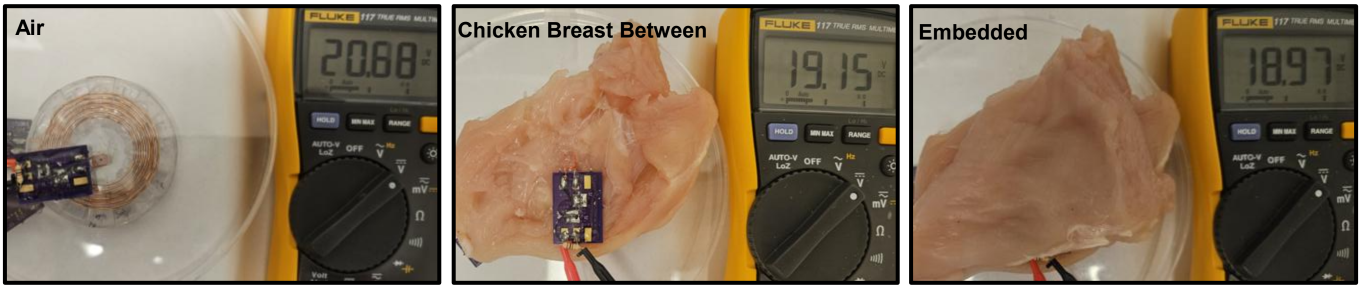


**Supplementary Fig 8. Tissue attenuation evaluation in air and chicken breast tissue.** The receiver was positioned between 1 cm thick layers of chicken breast tissue above and below the antenna (2 cm total tissue thickness). The measured voltages across a 1 kΩ load were 20.68 V in air, 19.15 V with the receiver placed beneath the chicken breast tissue, and 18.97 V with the receiver fully embedded between the upper and lower tissue layers.

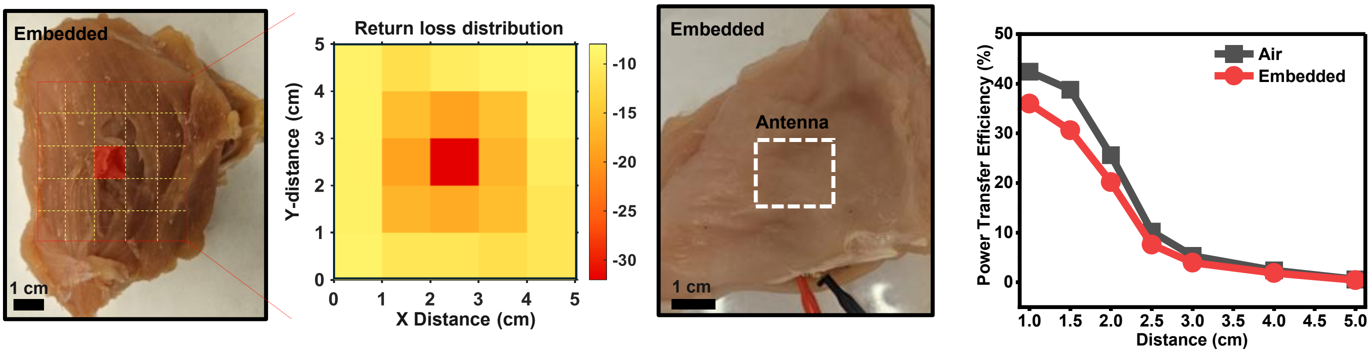


**Supplementary Fig 9. Localization evaluation in chicken breast tissue with the antenna embedded on the top and bottom surfaces.** Heat maps of the return loss and the received power as a function of distance within the chicken breast tissue are shown. The received power measured with the receiver embedded in the chicken breast tissue decreased by approximately 10% compared with that measured in air.

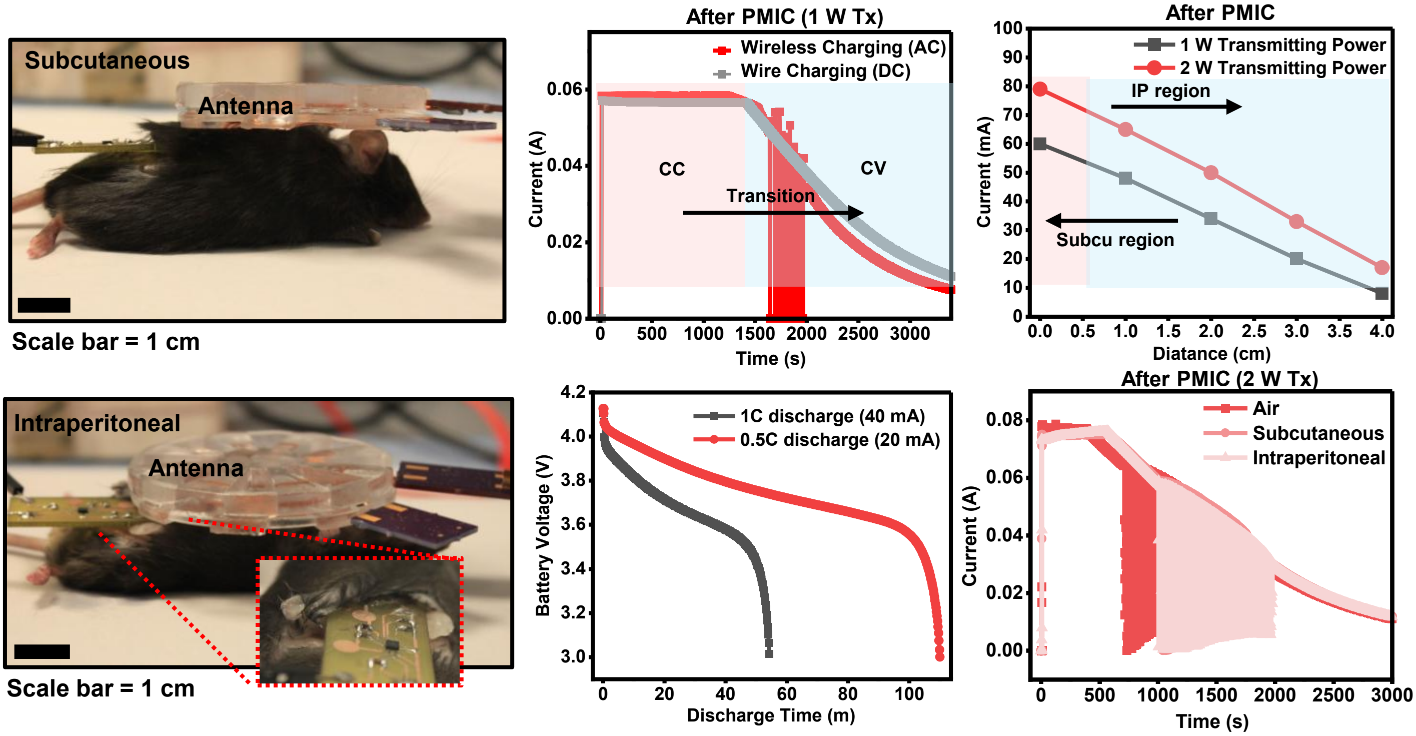


**Supplementary Fig 10. Charging of a 3.7 V, 40 mAh Li-ion battery.** Charging evaluation using a handheld transmitter and a millimeter-scale receiver implanted in the subcutaneous (SQ) and intraperitoneal (IP) spaces of cadaver mice. The charging current measured by direct DC charging under the same current condition as wireless charging is shown for comparison. The charging current was also measured at different transmitter (Tx)-to-receiver distances. To verify successful wireless charging, the charged battery was discharged under galvanostatic conditions. Charging curves obtained in air, the SQ space, and the IP space are compared, demonstrating similar charging profiles under all conditions.

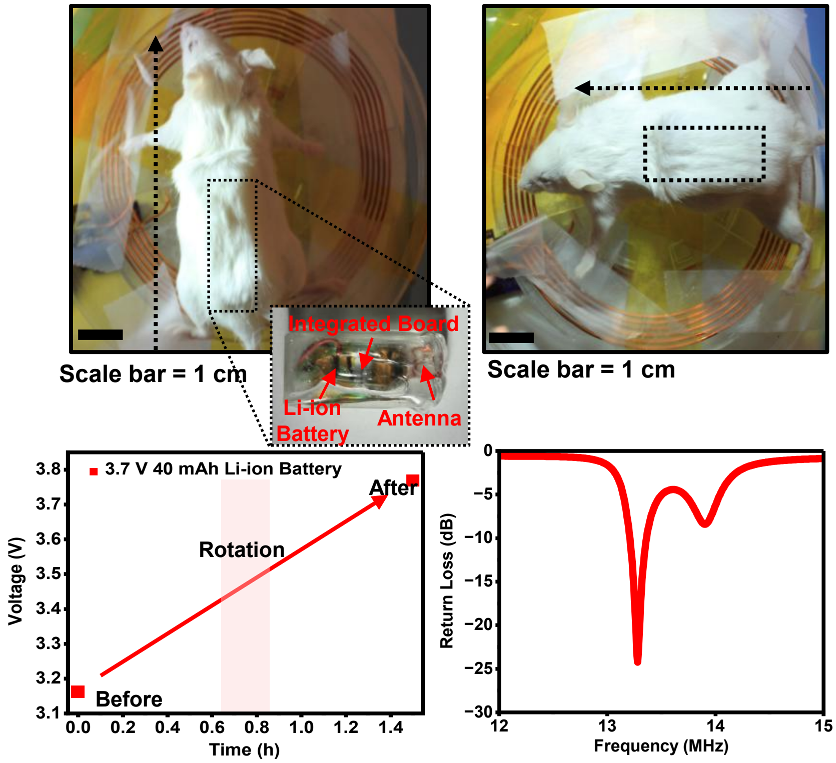


**Supplementary Fig 11.** Battery charging of the fully integrated device in cadaver mice using a larger transmitter antenna (Diameter = 10 cm). The fully integrated device, consisting of a PMIC, PCB, millimeter-scale antenna, 3.7 V, 40 mAh Li-ion battery, and encapsulation, is shown. The battery voltage before and after wireless charging is presented. During charging, the cadaver mouse implanted with the device was rotated at 45m to mimic a freely moving animal. The measured return loss of the fully integrated system is also shown for device localization.

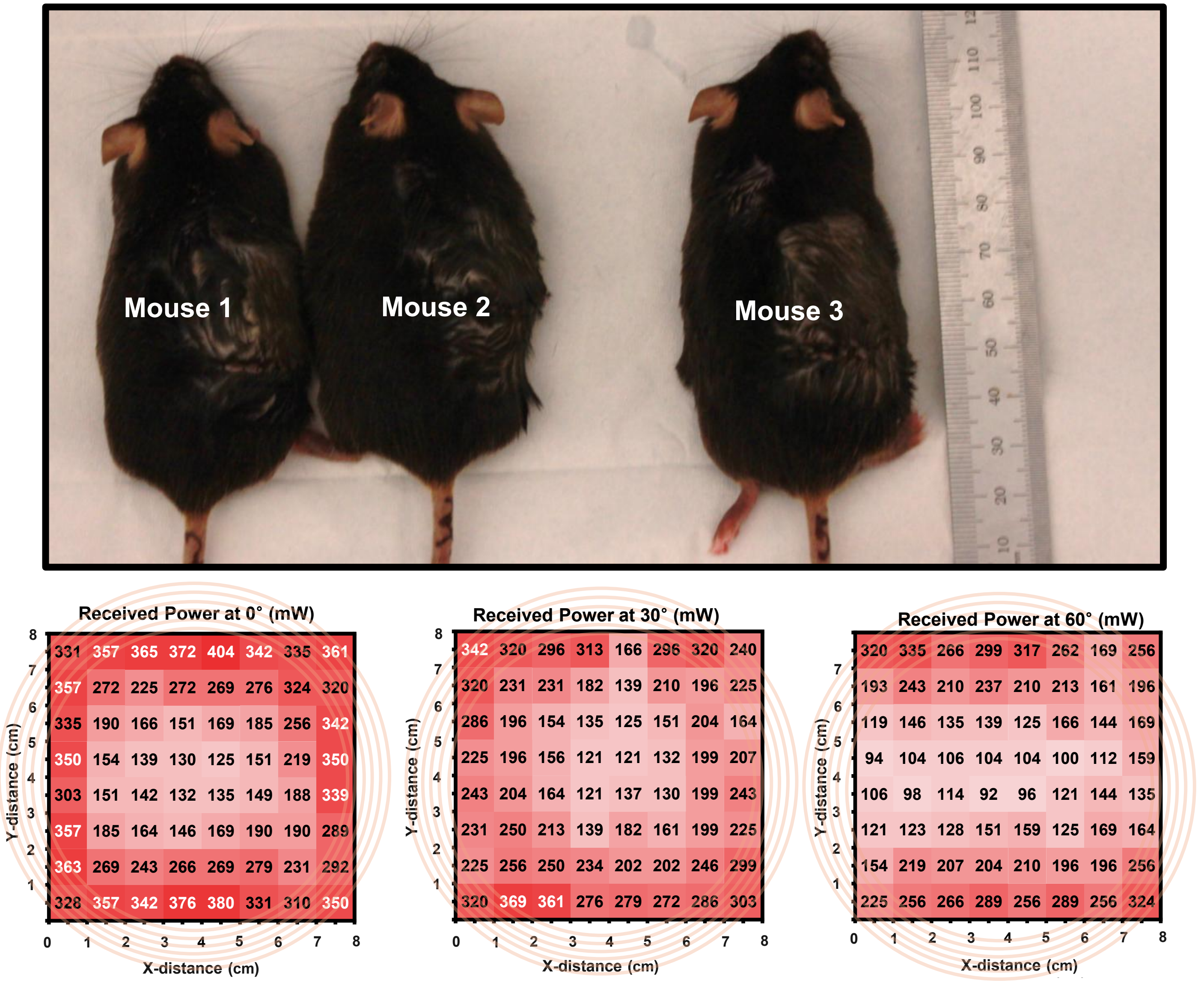


**Supplementary Fig 12. Freely moving mice charging.** Three fully integrated devices, each consisting of a 3.0 V, 5.8 mAh rechargeable coin-cell battery (ML621-TZ1, FDK America, Inc., Texas, USA), a power-management integrated circuit (PMIC), and an antenna, were implanted in three mice. The fully integrated device is shown alongside a U.S. dime and a Tylenol tablet for size comparison. The bottom row shows the received power measured in air at device orientations of 0°, 30°, and 60°, with a Rs–Rx separation of 1 cm.

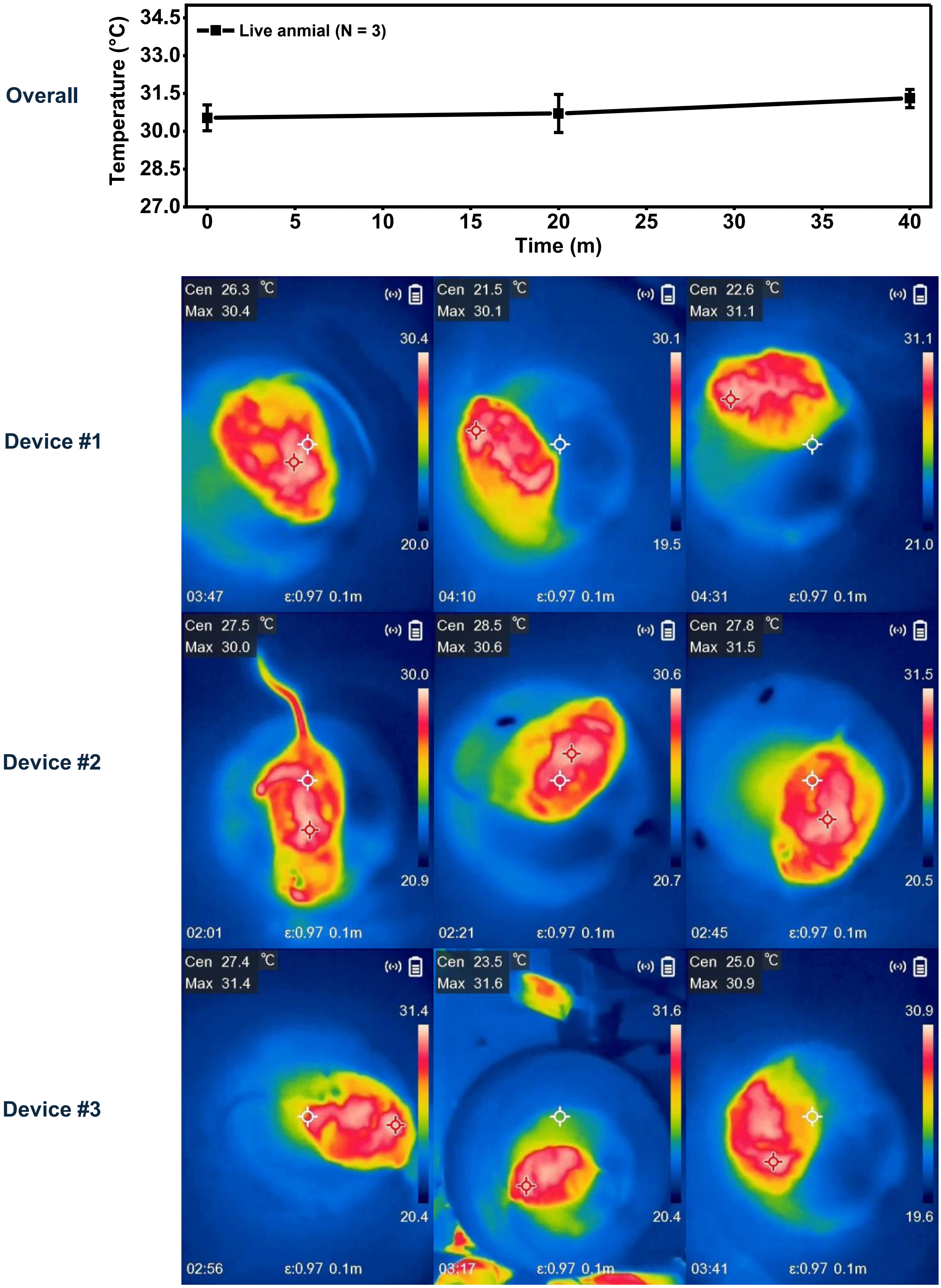


**Supplementary Fig 13. Temperature evaluation.** The temperature of freely moving mice during wireless power transfer was measured at three time points (0, 20, and 40 min). Thermal images were acquired using a near-infrared (NIR) camera to evaluate the thermal characteristics of the implanted integrated device. The average body temperature of freely moving mice during wireless power transfer measured at 0, 20, and 40 min was 30.53 ± 0.51 °C, 30.70 ± 0.75 °C, and 31.30 ± 0.36 °C, respectively (mean ± SD, n = 3).

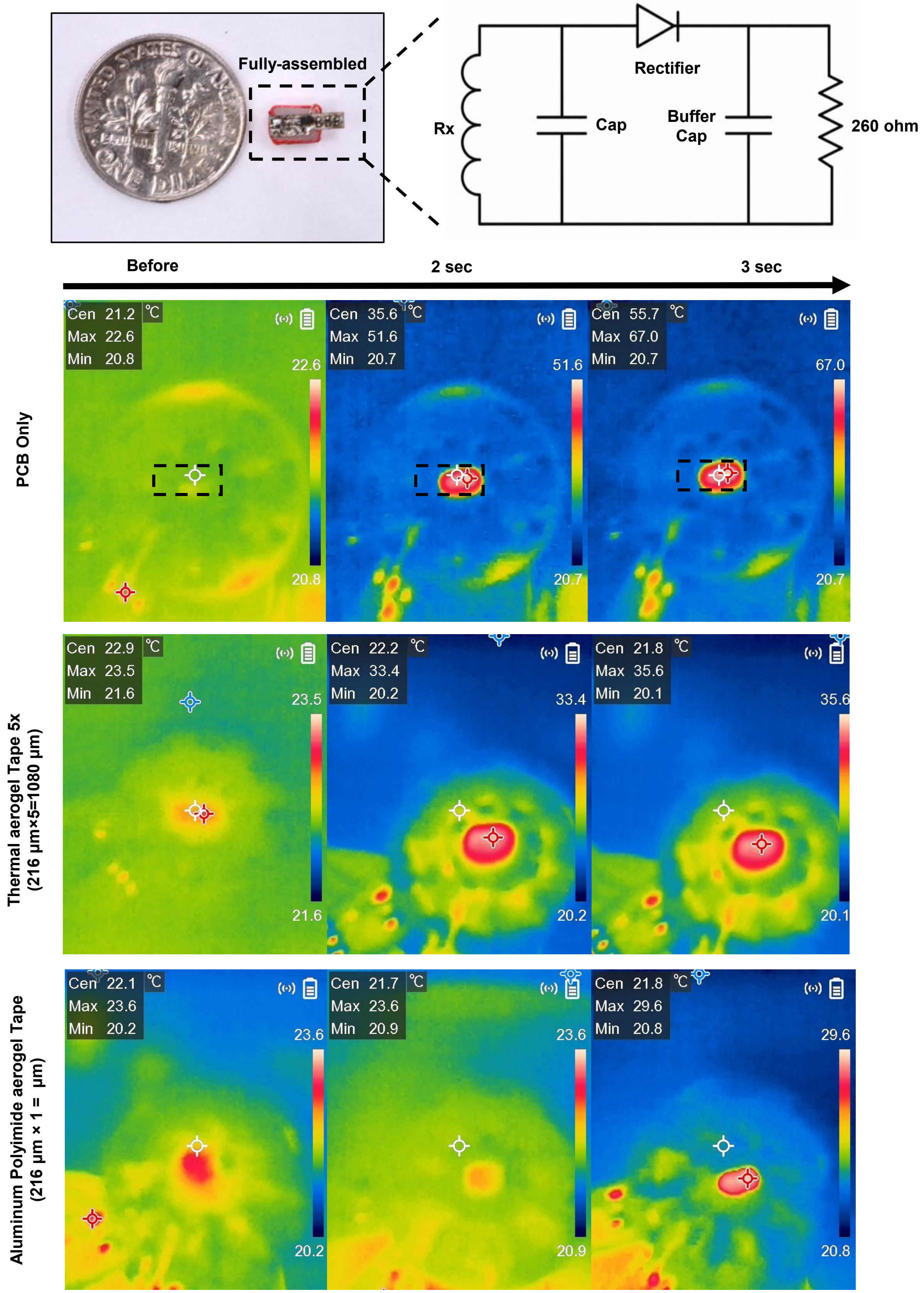


**Supplementary Fig 14.** Fully assembled device consisting of the antenna, printed circuit board (PCB), capacitors, diode, and resistors. The integrated device is shown alongside a U.S. dime for size comparison. The equivalent circuit of the device is illustrated on the right. The bottom panels shows near-infrared (NIR) thermal images acquired before and after wireless power delivery for thermal characterization. A localized temperature increase is observed only at the resistor, while no appreciable heating is detected in the surrounding components, confirming successful wireless power transfer, electrical power delivery, and localized thermal actuation. Furthermore, aerogel tape is attached to the fully assembled board with the x5 tape on top of the device.

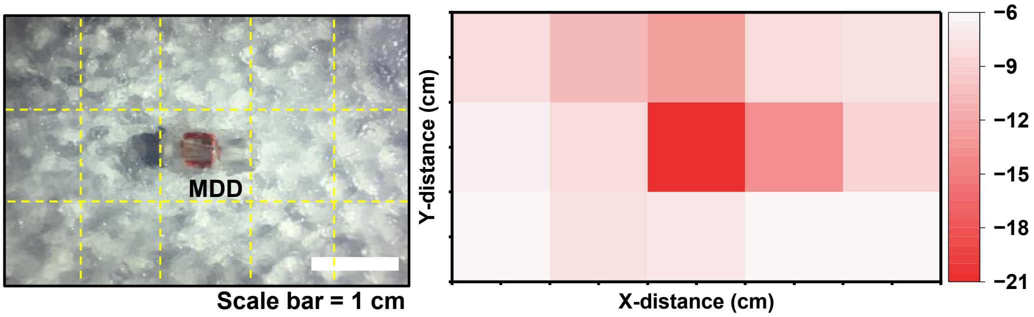


**Supplementary Fig 15.** Localization of the drug delivery device in 1% agarose gel. The spatial distribution of the return loss heat map is presented.

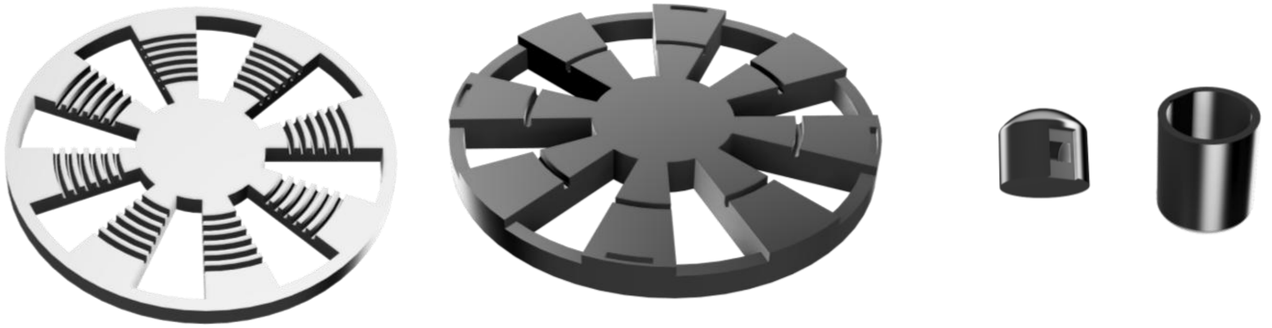

**Supplementary Fig 16.** All CAD files used in this work are presented. The left and middle images show the receiver (Rx) and transmitter (Tx), respectively. The right image shows the miniaturized drug delivery device encapsulation.

Without Saline

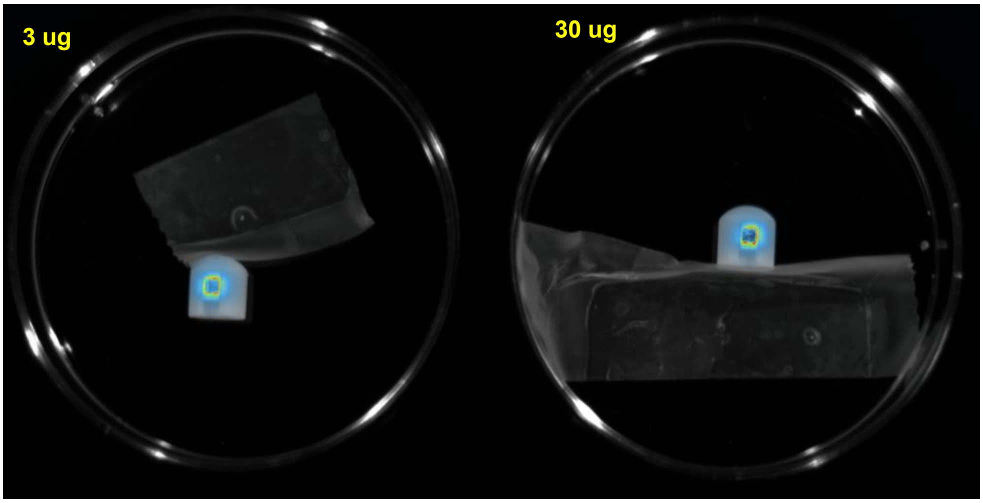


1 mL Saline

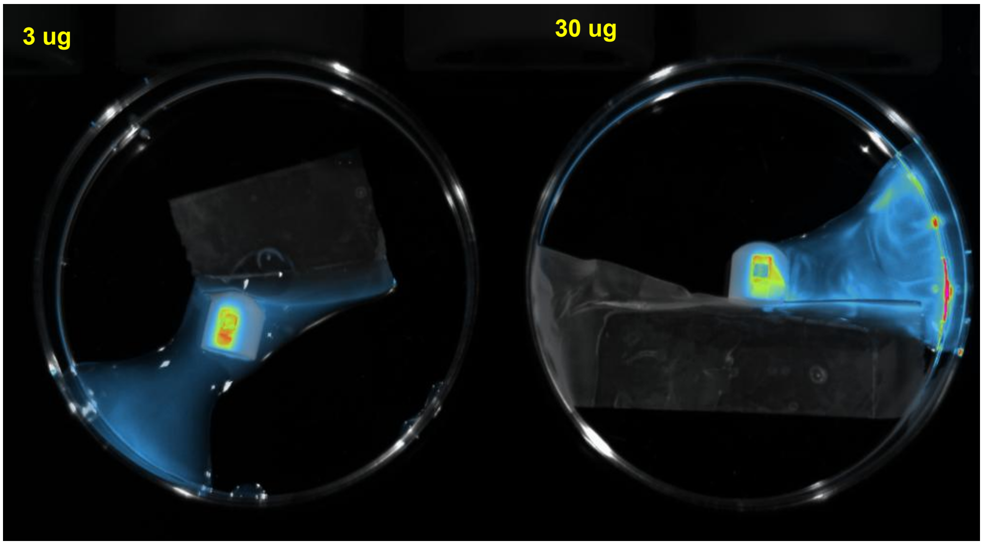


**Supplementary Fig 17.** Cy7 release in the drug release device chamber before and after adding saline under IVIS imaging**.**
