## Supplementary Note 1 for "Distributed resonant coupling enables high efficiency power transfer to mm-scale bioelectronics"

Corresponding author

Supplementary vidoes can be found in the following link;

<https://drive.google.com/drive/folders/133eaxrpLsp4To6ESQD9gjA6sfhGtIs3K?usp=sharing>

**Supplementary Note 1**

**First approximation of the DRC system**

To conceptualize the active participation mechanism in DRC, we hypothesize that the impedance of the spiral resonator is changed by the presence of the receiver (Rx). Specifically, maximum power transfer among the transmitter (Tx), resonator (Rs), and receiver (Rx) is achieved only when the Rx impedance actively modifies the impedance of the Tx–Rs system and drives the overall system into a strongly coupled resonant mode.

To test this hypothesis, we model the Tx, Rs, and Rx as three series RLC branches. Each branch *i* has an impedance given by

$$Z_{i}(\omega)=R_{i}+j\left( \omega L_{i} - \frac{1}{\omega C_{i}} \right),i\in\{T,S,R\}$$

(1)

where $R_{i}$, $L_{i}$, and $C_{i}$ are the resistance, inductance, and tuning capacitance of branch *i*, and the labels $t$, $s$, and $r$ denote the transmitter loop, the spiral resonator, and the receiver. Each branch is tuned so that $\omega L_{i}$equals $1/\omega C_{i}$ at the operating frequency $f_{0}$ of 13.56 MHz, at which point its reactance is cancelled and its impedance becomes optimal impedance to the loss resistance $R_{i}$.

To estimate the received voltage using a matrix formulation, we modeled the three lumped resonant elements as magnetically coupled resonators. The loop couples to the spiral through the mutual inductance $M_{TS}$, and the spiral couples to the receiver through $M_{SR}$. The direct coupling between the loop and the receiver is assumed to be negligible compared with the coupling path through the spiral resonator and is therefore neglected in this first-order approximation. This simplified model is intended to illustrate the underlying concept of DRC under the ideal condition for maximum power transfer. Applying Kirchhoff's voltage law to the three coupled branches, with the source voltage $V_{in}$ applied only to the loop, yields

$$Z_{T}I_{T}+j\omega M_{TS}I_{S}=V_{in}$$

(2a)

$$j\omega M_{TS}I_{T}+Z_{S}I_{S}+j\omega M_{SR}I_{R}=0$$

(2b)

$$j\omega M_{SR}I_{S}+Z_{R}I_{R}=0$$

(2c)

where $I_{T}$, $I_{S}$, and $I_{R}$are the branch currents.

**Receiver participation criterion by reflected impedance**

To calculate the effective impedance seen by the spiral resonator looking into the receiver (Rx), Eq. (2c) is rearranged to express the receiver current in terms of the spiral current:

$$\begin{matrix} & I_{R}=-\frac{j\omega M_{SR}}{Z_{R}}I_{S} & & \text{(3)} \end{matrix}$$

Substituting this into (2b) removes the receiver as an explicit variable and adds a single term to the Rs. With the consideration of impedance of the Tx seen by Rx is zero, the receiver therefore appears to the spiral as the reflected impedance

$$\begin{matrix} & Z_{R\to S}=\frac{\omega^{2}M_{SR}^{2}}{Z_{R}} & & \text{(4)} \end{matrix}$$

The Rx is reflected into the Rs as an equivalent impedance, $Z_{R\to S}=\omega^{2}M_{SR}^{2}/Z_{R}$. As a result, the Tx sees an effective Rs impedance of $Z_{S}+Z_{R\to S}$, which produces the reflected impedance $\omega^{2}M_{TS}^{2}/(Z_{S}+Z_{R\to S})$ at the Tx.

At the operating frequency, the reactive component of the receiver impedance is canceled, such that the receiver impedance reduces to an effective resistance, $R_{R}^{eff}$, which accounts for both the conductor loss and the reflected load. The reflected impedance in Eq. (4) therefore becomes

$$\begin{matrix} & \frac{\omega_{0}^{2}M_{SR}^{2}}{R_{R}^{eff}} & & \text{(5)} \end{matrix}$$

The Rx can influence the shared resonant mode only when the reflected resistance is sufficiently large to compete with the intrinsic loss of the spiral resonator. Therefore, the reflected loss should be comparable to or greater than the spiral loss, giving

$$\begin{matrix} & \omega_{0}^{2}M_{SR}^{2}\gtrsim R_{S}R_{R}^{eff} & & \text{(6)} \end{matrix}$$

Equation (6) can be rewritten in terms of the branch quality factors, $Q_{S}=\omega_{0}L_{S}/R_{S}$and $Q_{R}=\omega_{0}L_{R}/R_{R}^{eff}$, together with the coupling coefficient $k_{SR}$, where $M_{SR}=k_{SR}\sqrt{L_{S}L_{R}}$. Substituting these definitions into Eq. (6) yields the compact criterion

$$\begin{matrix} & k_{SR}\sqrt{Q_{S}Q_{R}}\gtrsim1 & & \text{(7)} \end{matrix}$$

Equation (7) is the three-element analogue of the strong-coupling criterion for two magnetically coupled resonators [1]. When this product exceeds threshold, the system enters the over-coupled regime, where the single resonant mode splits into two hybridized normal modes. In this regime, the receiver no longer behaves as a weakly coupled load but instead becomes part of the shared resonant state through active participation. Experimentally, the measured coupling-to-loss ratio of 1.9 and the observed mode splitting in the return-loss spectrum confirm that the millimeter-scale receiver operates in this over-coupled regime (Fig. 2b).

**Geometry-Dependent Criterion**

To derive a first order geometry-dependent design rule, we isolate the geometry-dependent contribution to the receiver impedance. Since the receiver electronics (rectifier and load) remain unchanged across different receiver designs, only the conductor resistance of the receiver coil varies with geometry. Therefore, the geometry-dependent contribution to the receiver impedance is approximated by the conductor resistance of the receiver coil ($R_{r}$)

$$\begin{matrix} & R_{r}=\rho\frac{\mathcal{l}}{A} & & \text{(8)} \end{matrix}$$

where $\rho$ is the resistivity of copper, $\mathcal{l}\approx Np$ is the total wire length for a receiver with $N$ turns and mean perimeter $p$, and $A=\pi d^{2}/4$ is the wire cross-sectional area for a wire diameter $d$. Therefore,

$$\begin{matrix} & R_{r}=\frac{4\rho N_{R}p}{\pi d^{2}} & & \text{(9)} \end{matrix}$$

For a fixed mean perimeter *p*, Eq. (9) shows that $R_{r}\propto\frac{N}{d^{2}}$. For a small receiver, the mutual inductance between the spiral resonator and the receiver can be approximated to first order as

$$\begin{matrix} & M_{SR}(z)=\frac{\mu_{0}N_{S}N_{R}A_{r}a_{S}^{2}}{2(a_{S}^{2}+z^{2})^{3/2}} & & \text{(10)} \end{matrix}$$

where $N_{S}$ and $a_{S}$ are the effective number of turns and radius of the spiral resonator, and $z$ is the separation between the spiral resonator and the receiver. For a fixed receiver capture area $A_{r}$, Eq. (10) shows that $M_{SR}\left( z \right)\propto N$, $M_{SR}^{2}\propto N^{2}$, whereas $R_{r}\propto N/d^{2}$. Therefore, substituting these scaling relationships into Eq. (6) yields Eq. (11), in which $M_{SR}^{2}\propto N^{2}$whereas $R_{r}\propto N/d^{2}$.

$$\begin{matrix} & \omega_{0}^{2}\left( \frac{\mu_{0}N_{S}N_{R}A_{r}a_{S}^{2}}{2(a_{S}^{2}+z^{2})^{3/2}} \right)^{2}\gtrsim R_{S}\frac{4\rho N_{R}p}{\pi d^{2}} & & \text{(11)} \end{matrix}$$

Therefore, one factor of $N$ remains after simplification, yielding the geometry-dependent scaling law in Eq. (12). After simplification,

$$\begin{matrix} & Nd^{2}\gtrsim K(a_{S}^{2}+z^{2})^{3}, & & \text{(12)} \end{matrix}$$

where $K$ is a constant that incorporates the operating frequency, material properties, and spiral resonator parameters. With this configuration, we can optimize the geometrical optimization of the Rx for the strongly-coupled region (Extended Fig 2.)

**Solution of the S11 splitting signature**

Assuming that the receiver (Rx) operates in the strong-coupling regime, the resulting eigenmodes are evaluated through the S-parameter response.

Solving Eq. (2) yields the three branch currents and, consequently, all measurable electrical quantities. The input impedance at the transmitter is

$$\begin{matrix} & Z_{in}=Z_{T}+\frac{\omega^{2}M_{TS}^{2}}{Z_{S}+\omega^{2}M_{SR}^{2}/Z_{R}} & & \text{(13)} \end{matrix}$$

from which the reflection coefficient measured by the vector network analyzer is

$$\begin{matrix} & S_{11}=\frac{Z_{in}-Z_{0}}{Z_{in}+Z_{0}} & & \text{(14)} \end{matrix}$$

where $Z_{0}$ is the reference impedance.

Because the Rx is reflected into the Rs through magnetic coupling, the resonance of the system is no longer determined by the Rs alone. Instead, the coupled Rs-Rx subsystem behaves as a single hybrid resonator whose resonance frequencies depend on both the spiral impedance and the reflected receiver impedance. Consequently, the reflection minima correspond to the natural eigenmodes of the coupled resonator pair rather than the isolated resonance of either element.

Once the coupled mode is established, electromagnetic energy oscillates coherently between the Rs and the Rx, rather than being transferred only once to the receiver. This coherent energy exchange allows the receiver to actively participate in the shared resonant mode, resulting in the formation of hybridized eigenmodes. The reflection minima occur at the natural resonances of the coupled Rs–Rx system, corresponding to the frequencies at which

$$\begin{matrix} & Z_{S}(\omega)Z_{R}(\omega)+\omega^{2}M_{SR}^{2}=0 & & \text{(15)} \end{matrix}$$

Unlike an independent resonator, Eq. (15) presents two eigen-frequency solutions when the magnetic coupling is sufficiently strong. The appearance of two resonance minima therefore provides an experimental signature that the Rx is no longer a passive load but has become an active participant in the shared resonant mode.

Physically, the electromagnetic energy oscillates between the Rs and the Rx, producing two hybridized resonant modes with an in-phase mode and an out-of-phase mode. This mode hybridization lifts the degeneracy of the original resonance and produces the characteristic resonance splitting observed in the return-loss. This characteristic equation has two solutions, giving rise to the observed resonance splitting. In the low-loss limit, with both resonators tuned to $\omega_{0}$, the resonance frequencies are

$$\begin{matrix} & f_{\pm}=\frac{f_{0}}{\sqrt{1\mp k_{SR}}}, \Delta f\approx k_{SR}f_{0} & & \text{(16)} \end{matrix}$$

predicting the mode splitting observed experimentally. As the coupling becomes stronger, the two eigenmodes separate further in frequency, resulting in the wider peak splitting observed experimentally. Conversely, when the coupling is weak, the two modes collapse into a single resonance and the splitting disappears.

The resonance frequencies are determined primarily by Eq. (16), whereas the depth of each reflection minimum depends on the impedance transformation between the coupled modes and the reference impedance $Z_{0}$.

For the practical device, which includes loss, detuning, and geometric variations represented by the configuration parameter $\xi$, the resonance pair is determined numerically as the two local minima of the modeled reflection in sweep frequencies,

$$\begin{matrix} & f_{-}^{pred}(\xi),\text{ }f_{+}^{pred}(\xi)=\underset{f}{\mathrm{locmin}}\mid S_{11}(2\pi f,\xi)\mid& & \text{(17)} \end{matrix}$$
